# Sulindac modulates the response of triple negative breast cancer to anti-PD-L1 immunotherapy

**DOI:** 10.1101/2025.06.11.659159

**Authors:** Bin Yi, Ruixia Ma, Xin Zhang, Li Li, Adam I. Riker, Yaguang Xi

## Abstract

Teriple-negative breast cancer (TNBC) is a heterogeneous tumor, and there is a lack of effective therapies. Immune checkpoint inhibitors (ICIs) therapy has been widely used to treat a variety of human cancers including TNBC. Despite the FDA’s approval of therapies like Atezolizumab (anti– PD-L1) and Pembrolizumab (anti-PD-1) for certain TNBC cases, more than half of all TNBC patients, especially those with low PD-L1 levels and in advanced stages, remain unresponsive. Therefore, there is urgent clinical need to find other innovative combination therapy strategy for these TNBC patients. In this study, we examined the efficacy of sulindac to enhance the response of TNBC to anti-PD-L1 immunotherapy. We utilized a 4T1 syngeneic mouse tumor model to compare the inhibitory effects of PD-L1 antibody, sulindac, and their combination on 4T1 tumor growth. We found that mice treated with combination therapy showed a significant reduction in tumor volume, along with increased infiltration of activated T lymphocytes (Granzyme B+/CD8+ T cells) in the tumor tissues. We also established a PBMC humanized mouse model to further confirmed that combination therapy could significantly reduce the tumor size of TNBC patient-derived xenografts (56S) and the combination group were found to have the most infiltrating activated human T lymphocytes (Granzyme B+/CD8+ T cells) in the tumor tissues. Immunofluorescent staining of organoids also confirmed that organoids from combination group have more CD8+ T cell and the results of organoid cell viability assay *in vitro* indicated that combination of sulindac with PD-L1 antibody together with activated human PBMC significantly reduce 56S organoid cell viability, which is consistent with *in vivo* study. When investigating the mechanism of action, first we found sulindac could downregulate exosomal PD-L1 by decreasing expression of nsMase2 which is a major regulator in production of exosomal PD-L1. Second, we demonstrated that sulindac could downregulate PD-L1 by blocking Stat3 signaling and enhancing the expression of miR-570-3p which can potentially target PD-L1, which in turn led to a further decrease in exosomal PD-L1. Previous study showed that PD-L1 antibody could be bound and consumed by exosomal PD-L1 in the blood circulation. Therefore, in combination therapy, sulindac downregulating exosomal PD-L1 leads to increased availability of PD-L1 Ab, which potentially improves the overall efficacy of anti-PD-L1 therapy. In conclusion, our findings provide unique insights into the mechanism of action and efficacy for sulindac as an immunomodulatory agent in combination with anti-PD-L1 therapy for the treatment of TNBC.

## Introduction

Currently, breast cancer remains the second leading cause of death among women in the United States, with an estimated 43,250 deaths in 2022 [1]. Triple-negative breast cancer (TNBC), which lacks significant expression of estrogen receptor (ER), progesterone receptor (PR) and human epidermal growth factor receptor 2 (HER 2), accounts for approximately 20% of all breast cancers [2]. Epidemiological studies have reported African-American women with TNBC have worse clinical outcomes compared with European women with TNBC, and this disease is more prevalent in poor socioeconomic groups [3]. Because TNBC has aggressive phenotypes including a high prevalence in young women with breast cancer, early recurrence, and distant metastasis, TNBC patients have a high rate of poor prognosis [4–6]. Through analyzing 1,601 patients with breast cancer, one group found that TNBC patients showed an increased risk of distant recurrence compared with non-TNBC patients and TNBC patients have a high risk of distant recurrence during the first 3 years after therapy [5]. Lacking ER, PR and HER2 expression, TNBC is a problematic and unpredictable malignancy and there is no effective treatment for TNBC patients. As such, there is an urgent medical need to develop more effective and safer treatment options for TNBC patients.

Recently, Immune checkpoint inhibitors, specifically PD-1 and PD-L1, have demonstrated significant inhibitory efficacy against various cancers [7, 8] The PD-1 cell receptor, primarily expressed on immune cells such as T cells and macrophages [9, 10], is a T cell co-suppressor receptor. In contrast, its ligand, PD-L1, is mainly expressed on dendritic cells (DCs) and various types of tumor cells. Under normal physiological conditions in humans, the interaction between PD-1 and PD-L1 appears to protect against the development of autoimmune diseases [11]. However, when activated T lymphocytes recognize tumor cells, elevated PD-L1 levels on the tumor cell surface can bind to PD-1 on T lymphocytes. This interaction inhibits the tumor-killing ability of T lymphocytes, leading to immune surveillance failure [12]. Consequently, the fundamental principle of immune checkpoint inhibitor (ICI) immunotherapy is to block the direct binding between PD-1 and PD-L1, thereby restoring the surveillance function and tumor-killing capacity of T lymphocytes on tumor cells [13].

Immune checkpoint inhibitors (ICIs) therapy has been widely used to treat a variety of human cancers including TNBC. Several clinical studies have shown that PD1/PD-L1 monotherapy demonstrated modest response to advanced mTNBC with positive PD-L1 [6, 14–16] which suggests that combination therapy may improve the outcome of ICIs immunotherapy against TNBC. The FDA approved atezolizumab (anti–PD-L1) in combination with nab-paclitaxel for the treatment of patients with unresectable locally advanced or metastatic PD-L1 positive TNBC in 2019 and granted accelerated approval of Pembrolizumab (anti-PD-1) in combination with chemotherapy for patients with locally recurrent unresectable and metastatic PD-L1 high positive TNBC in 2020. Then, FDA approved pembrolizumab for high-risk early-stage triple-negative breast cancer in combination with chemotherapy as neoadjuvant treatment, and then continued as a single agent as adjuvant treatment after surgery in 2021. Unfortunately, such combination strategies just benefit a small group of TNBC patients with high levels of PD-L1. Therefore, there is an urgent clinical need to find other innovative combination therapy strategies for these TNBC patients.

We have long studied the chemopreventive effects of sulindac on TNBC, an FDA-approved nonsteroidal anti-inflammatory drug (NSAID) that has been shown to inhibit the development and progression of TNBC [17, 18]. In this study, we treated 4T1 syngeneic mice and 56S PDX PBMC humanized mice with low doses of sulindac in combination with PD-L1 antibody. Our results showed that two mouse models treated with the combination therapy [sulindac + PD-L1 antibody] demonstrated an improved response to tumor progression, compared to monotherapy PD-L1 antibody or sulindac alone. Of interest, infiltrating CD8+ T-cell especially the activated CD8+ T cells were significantly increased in tumor tissues of mice treated with combination therapy and combination therapy improved the killing effect on 56S tumor organoid. We demonstrate the mechanism of action is that sulindac can downregulate exosomal PD-L1 by decreasing expression of nsMase2 which is a key regulator of exosome PD-L1 production, blocking Stat3 signaling, and enhancing the expression of miR-570-3p which can potentially target PD-L1. Thus, our findings provide evidence that sulindac acts as an immunomodulator to enhance the response of TNBC tumors to anti-PD-L1 therapy.

## Materials and Methods

### Cell culture and reagents

The murine breast cancer cell line 4T1 and E0771 and the human breast cancer cell lines MDA-MB-231, HCC70 and BT549 were purchased from ATCC (Manassas VA, USA). Cell culture media were purchased from Thermo Fisher Scientific and mixed with 10% fetal bovine serum for cell culture. Cells were cultured at 37°C and 5% CO_2_ in a humidified incubator. Sulindac and CMC (carboxymethylcellulose) were purchased from Sigma-Aldrich (St Louis MO, USA). CMC was used as the vehicle control of sulindac for *in vivo* study. Stat3 pcDNA3 vector and Stat3 inhibitor (Stattic) are gifts from Dr.Qiang Sheng. Anti–mouse PD-L1 antibody and isotype control rat IgG2b were purchased from Bio X Cell (West Lebanon NH, USA). Anti-human PD-L1 antibody (Atezolizumab) and human normal PBMCs isolated from healthy adult donors were from Ochsner health. Sulindac sulfide was purchased from Alfa Aesar (Haverhill MA, USA).

### Animal study

BALB/c immunocompetent mice and NOD/SCID immunodeficiency mice were purchased from Charles River Laboratories (Wilmington MA, USA). The animals were housed in specific pathogen-free conditions, and all animal experiments were performed in accordance with protocols approved by the Institutional Animal Care and Use Committee of the Louisiana State University Health Sciences Center.

4T1 syngeneic mouse model with mammary fat pat injection and PBMC humanized mouse model were established. In brief, BALB/c mice were first anesthetized with vaporized isoflurane and oxygen for 4T1 syngeneic mouse model. The incision site was then shaved and disinfected with iodine. Afterwards, a small incision was made, and 1×10^5^ 4T1 tumor cells were subcutaneously implanted in the mammary fat pad of each mouse. When 4T1 tumors reach about 50 mm^3^, which was 7 days after tumor cell injection, 28 female BALB/c mice were randomly divided into 4 groups (7 mice per group): (1) vehicle control treated with CMC + IgG2b; (2) sulindac (7.5 mg/kg, p.o., bid.); (3) anti-mouse PD-L1 antibody (50 μg, i.p. b.i.w.); (4) combination of sulindac and anti-mouse PD-L1 antibody (7.5 mg/kg p.o., bid. + 50 μg, i.p. b.i.w.). For PBMC humanized mouse model, NOD/SCID mice were anesthetized with vaporized isoflurane, and the incision site was shaved and disinfected with iodine, as described for the 4T1 model. Then 56S patient-derived xenograft (PDX) tumor (approximately 5 x 4 mm pieces) were implanted into mammary fat pat of each mouse. The incision was closed using non-absorbable sutures, which were removed after the wound had healed. Two independent 56S animal experiments were performed. When 56S tumors reached about 70 mm^3^, mice were pretreated with sulindac for up to 1 week, and 2×10^7^ human PBMCs were intraperitoneally injected into each mouse. For the first experiment, twelve female NOD/SCID mice were randomly divided into four groups (3 mice per group), and for the second experiment, twenty four female NOD/SCID mice were randomly divided into four groups (6 mice per group): (i) vehicle control treated with CMC; (ii) sulindac (7.5 mg/kg, p.o., b.i.d.); (iii) anti-human PD-L1 antibody (200 μg, i.v. q.week); (iv) combination of sulindac and anti-human PD-L1 antibody (7.5 mg/kg p.o., b.i.d. + 200 μg, i.v. q.week).

Mice were monitored for tumor growth bi-weekly using external caliper measurements (tumor volume= L×W^2^/2, where L represents the tumor length and W represents the tumor width) until the endpoint. BALB/c mice were euthanized on day 27 after 4T1 cells injection, and NOD/SCID mice were euthanized after human PBMCs injection for up to 4 weeks. Blood samples were collected for exosome isolation, tumor tissue samples were fixed in 10% buffered formalin for paraffin embedding, and tumor organoid were isolated for CD8+ T cell staining and tumor organoid viability assay.

### Isolation and culture of tumor organoids

The 56S tumor organoids were isolated and cultured for CD8+ T cell staining and tumor organoid viability assay. Briefly, in a cell culture hood, 56S tumors were placed in a 6 cm dish with 1 mg/ml of type IV collagenase medium (Thermo Fisher Scientific, Worcester MA, USA), and the tumors were cut into small pieces with a sterile knife. These small tumor pieces were then transferred to collagenase medium tubes and incubated in a 37°C water bath, and then we need to check for right formation of spheroid every 5-10 minutes. The same volume of RPMI 1640 medium containing 5% FBS was added to the collagenase medium tube to neutralize the collagenase digestion. The collagenase medium tube was then spined down at RT 500 × g for 5 min and the supernatant was discarded to obtain minced tumor. After passing the minced tumor over a 100 µm filter (VWR, Radnor, PA, USA) placed on top of a 50 ml tube (S1 fraction) and smashing the tumor chunks retained on the filter with the plunger of syringe, the filter is then washed with RPMI 1640 medium containing 5% FBS thereby obtaining the S1 fraction. After passing the S1 fraction over a 70 µm filter (VWR) placed on top of a 50 ml tube, the 70 µm filter was placed upside down on top of a new 50 ml tube and washed with RPMI 1640 medium containing 5% FBS to obtain the S2 fraction. The 56S tumor organoids were collected after spinning down the S2 fraction at 500 × g for 5 min and removing the supernatant. The 56S tumor organoids were resuspended in collagen (Thermo Fisher Scientific) (total=500 µl including 50 µl 10x PBS, 417 µl 3 mg/ml Collagen, 12.5 µl 0.5 M NaOH and 20.5 µl sterile H_2_O) and plate 75 µl per well in an 8-well chamber (Fitchburg, WI, USA) for the following CD8+ T cell staining.

### Immunofluorescence staining

Immunofluorescence staining was performed on 56S tumor organoid, paraffin-embedded sections of 4T1 and 56S tumor tissue and TNBC cell lines (4T1, E0771, MDA-MB-231 and HCC70). The paraffin-embedded sections were deparaffinized and rehydrated. After antigen retrieval and permeabilization, slides were blocked with 5% horse serum for 1 hour at room temperature and then incubated with the primary antibodies at 4°C overnight. To stain CD8+ T cells in 4T1 tumor tissues, tumor tissue slides were incubated with Alexa Fluor® 647–conjugated CD8a antibody (Biolegend, San Diego CA, USA) overnight. To stain CD8+ T cells in 56S tumor tissues, tumor tissue slides were incubated with FITC mouse anti-human CD8 antibody (BD Bioscience, Sparks, MD, USA) overnight. For CD8+ T cells staining in the 56S tumor organoids from 4 *in vivo* groups, the organoids were incubated with FITC mouse anti-human CD8 antibody (BD Bioscience) for overnight at 37°C and 5% CO2 in a humidified incubator, followed by 5 µl/ml of Hoechst 33342 solution (BD Bioscience) for 30 min. For CD8+ T cells staining in the 56S control tumor organoids mixed with 1×10^6^ activated human PBMCs, the tumor organoids were treated with vehicle control DMSO at 0.1%, 20 µM sulindac sulfide (SS), 200 nM anti-human PD-L1 antibody (Atezolizumab), and combination of 20 µM SS with 200 nM anti-human PD-L1 antibody (Atezolizumab) for 2 days. The 56S tumor organoids were then incubated with FITC mouse anti-human CD8 (BD Bioscience) for overnight at 37°C and 5% CO_2_ in a humidified incubator, followed by 5 µ/ml of Hoechst 33342 solution (BD Bioscience) for 30 min. For PD-L1 and Phospho-Stat3 staining, tumor tissue slides were incubated with primary mouse PD-L1 antibody (Abcam, Cambridge MA, USA) or human PD-L1 antibody (Cell Signaling, Beverly MA, USA) and primary Phospho-Stat3 antibody (Cell Signaling), followed by Alexa Fluor® 488 and 555–conjugated secondary antibody (Thermo Fisher Scientific) for 1 hour at room temperature, respectively. For immunofluorescence staining of all four TNBC cell lines, cells were seeded in flow dishes at 37°C overnight and then treated with 10 µM Stat3 inhibitor (Stattic) for 2 hours and 20 µM sulindac sulfide (SS) for 12 hours. DMSO at 0.1% was used as vehicle control. Mouse IL6 (VWR) and human IL6 (VWR) were added to the cells at a concentration of 50 ng/mL and incubated for 30 minutes, respectively. Subsequently, cells were fixed by 4% formaldehyde (Alfa Aesar) for 10 minutes, followed by permeabilization using 1% Triton X-100 (Bio-Rad, Hercules, CA, USA). After blocking with 1% BSA, cells were incubated with mouse PD-L1 antibody (Abcam), human PD-L1 antibody (Cell Signaling), and Phospho-Stat3 antibody (Cell Signaling) at 4°C overnight. After washing with PBS, cells were incubated with Alexa Fluor 488 or 555–conjugated secondary antibody (Thermo Fisher Scientific) for 1 hour at room temperature. Finally, after washing and staining with 5 ng/mL DAPI (Sigma-Aldrich), slides and cells were processed with Prolong Diamond Anti-fade Mountant (Thermo Fisher Scientific) and analyzed by confocal microscopy (Nikon Eclipse Ti2).

### Flow cytometry analysis

Referring to a published protocol [19], we analyzed T cell status in tumors from BALB/c and NOD/SCID mice using flow cytometry. First, we prepared single-cell suspensions from the tumor of BALB/c and NOD/SCID mice through mechanical dissociation. Briefly, after removing fat, necrotic tumor tissue and blood, the tumor tissues were washed in warm DMEM and then cut into 1-3 mm^3^ pieces. The tumor pieces were then minced and passed through a 40 µm cell strainer (Thermo Fisher Scientific) to remove clumps. The filtrate containing the single-cell suspensions was washed with DMEM and PBS, then centrifuged at 300 × g for 10 minutes at room temperature. T cells from normal human PBMCs and normal mouse blood were used as positive controls. For human PBMCs activation, we seeded 1 × 10^6^ human PBMC cells per well in a 24-well plate and then treated them with 1 μg/mL anti-human CD3 antibody (BD Bioscience) and 1 μg/mL anti-human CD28 antibody (BD Bioscience). The cell suspensions were counted and blocked with human or mouse Fc-blocked antibody (BD Biosciences) at 4°C prior to incubation with live-dead fixable viability stain 450 (BD Bioscience) for 20 min and flow cytometry antibodies for 30 minutes. These antibodies included PerCP-Cy5.5 anti-human CD45 antibody (BD Biosciences), APC-Cy7 anti-human CD3 antibody (BD Biosciences), APC anti-human CD8 antibody (BD Biosciences), BV711 anti-human HLA antibody (BD Biosciences), and PE-CF594 anti-human CD4 antibody (BD Biosciences) or APC anti-mouse CD45 antibody (BD Bioscience), BV711 anti-mouse CD3e antibody (BD Bioscience), PE-CF594 anti-mouse CD8a antibody (BD Bioscience), and APC-Cy™7 anti-mouse CD4 antibody (BD Bioscience).Subsequently, intracellular staining was performed using the BD GolgiStop (BD Biosciences) fixation and permeabilization solution kit. FITC anti-human Granzyme B antibody (BD Biosciences) or FITC anti-mouse granzyme B antibody (eBioscience) was incubated with the cells for 30 minutes. After washing the cells once with PBS, they were fixed with 1% paraformaldehyde and then subjected to flow cytometric analysis. Flow cytometry was performed by Beckman coulter Gallios and data were analyzed by Kaluza software.

### Tumor organoid viability assay

Referring to a published protocol [20], we analyzed tumor organoid viability after treatment with sulindac sulfide and anti-human PD-L1 antibody, with addition of either non-activated or activated human PBMCs. Tumor organoid viability was assessed using Cell Titer-Glo Assay (Promega, Madison, WI, USA), which measures viable cells based on quantification of ATP contents. Briefly, 32 μl of 7.5 mg/ml BME-2 (Cultrex® Reduced Growth Factor Basement Membrane Extract, Type 2, PathClear) (R&D Systems, Minneapolis, MN, USA) was dispensed into wells of a 96-well flat clear bottom polypropylene plate. After centrifugation at 300 × g for 1 minute to ensure BME-2 coverage at the bottom of each well, the 96 well plate was placed in a 37°C, 5% CO2 incubator for 30 minutes to facilitate polymerization. 56S tumor organoid embedded within 50 µl of organoid media (RPMI 1640 medium containing 5% FBS) including 1 µl of 7.5 mg/ml BME-2 were seeded in 96-well plates at a density of 1,000 organoids per well and incubated for 12 hours before treatments. Subsequently, 56S tumor organoids were treated with 0.1% DMSO vehicle control, 20 µM sulindac sulfide (SS), 200 nM anti-human PD-L1 antibody (Atezolizumab), and a combination of 20 µM SS with 200 nM anti-human PD-L1 antibody (Atezolizumab) and incubated for an additional 72 hours. At the end of the incubation, the relative cell viability was computed and analyzed by using GraphPad Prism 7 (GraphPad Software, CA, USA).

### Isolation and purification of exosomes from mouse plasma samples and cell culture media

Exosomes from mouse plasma samples were isolated and purified with the ExoQuick ULTRA EV isolation kit (System Biosciences, Mountain View, CA, USA). After pretreatment with 1 μL of Thrombin to remove fibrin, 125 μL of plasma was incubated with 33.5 μL of ExoQuick exosome precipitation solution for 30 minutes at 4°C, following the manufacturer’s instruction. After centrifuging the ExoQuick/plasma at 3,000 × g for 10 minutes, the exosomes pellet was enriched at the bottom of the tubes and then resuspended with buffer in the kit. Exosome from cell culture media were isolated and purified with ExoQuick-TC PLUS Exosome Purification Kit (System Biosciences). Exosome concentrations were measured using the NanoSight NS300 (Malvern Panalytical, Malvern, Worcestershire, United Kingdom). Lastly, exosome lysates were generated using RIPA buffer (Thermo Fisher Scientific) and analyzed by Western blot.

### Western blot assay

Total proteins lysates were prepared with RIPA buffer and quantified with the DC Protein Assay kit (Bio-Rad). After the denatured proteins were separated on a 10% SDS-PAGE gel, they were transferred to a PVDF filter (Bio-Rad). After blocking with 5% non-fat milk-TBST (Bio-Rad), the blots were incubated mouse PD-L1 antibody (Abcam), human PD-L1 antibody (Cell Signaling), β-actin antibody (Cell Signaling), mouse CD63 polyclonal antibody (Abcam), human CD9 antibody (Cell Signaling), nsMase2 antibody (Abcam), mouse Calnexin polyclonal antibody (Abcam) or human Calnexin polyclonal antibody (Abcam) at 4°C overnight, respectively. After washing three times with 1× TBST (Bio-Rad), the blots were incubated with peroxidase-linked secondary goat anti-rabbit IgG Ab (Bio-Rad), or goat-anti-mouse IgG Ab (Bio-Rad) for 1 hour at room temperature. Finally, visual imaging was conducted by VersaDoc Imaging System (Bio-Rad) after incubation with SuperSignal West Pico PLUS chemiluminescent substrate (Thermo Fisher Scientific). Volume One software (Bio-Rad) was utilized for the relative quantification of blotted proteins.

### Real-time PCR

Total RNA was extracted by TRIzol® (Thermo Fisher Scientific) and cDNA was synthesized by a high-capacity cDNA reverse transcriptase kit (Thermo Fisher Scientific). The stem-loop RT primers of miRNAs were designed following the previous publication [21]. The 20 μl mixture of reverse transcription reaction included 2 μg of total RNA, 2 μM reverse transcription primers, 2 μl 10× reverse transcription buffer, 0.8 μl 100 mM dNTP and nuclease-free water. The reverse transcription reaction was performed at 37°C for 2 h. The 20 μl mixture of quantitative real-time PCR reaction contains 10 μl 2× SYBR master mix (Thermo Fisher Scientific), 1 μl forward primer (7 μM) and 1 μl reverse primer (7 μM), 1 μl cDNA and 7 μl nuclease-free water. The real-time PCR was performed for 30 cycles on a QuantStudio 3 Real-Time PCR System (Thermo Fisher Scientific). Each cycle of real-time PCR includes denaturing for 10 s at 94°C, annealing and extension for 30 s at 58°C. The comparative Ct method was performed to analyze the relative expression of target miRNAs [21].

### Enzyme-linked immunosorbent assay (ELISA)

Mouse plasma PGE2 levels were measured with the Prostaglandin E2 Parameter Assay Kit (R&D Systems) according to the manufacturer’s instructions. Briefly, 50 μL of appropriately diluted plasma samples were added to a 96-well polystyrene microplate coated with goat anti-mouse polyclonal antibodies. After incubation with 50 μL of mouse PGE2 monoclonal antibody for 1 hour at room temperature, PGE2 conjugate was added to each well and incubated for 2 hours with gentle shaking. Following washing, the substrate solution was added to each well and incubated for 30 minutes before adding the stop buffer to halt the color development reaction. Finally, absorbance was measured at 450 nm using a microplate reader (Bio-Rad).

### Chromatin immunoprecipitation assay (ChIP)

The EZ-Magna ChIP kit (MilliporeSigma, Burlington, Massachusetts, USA) was used for ChIP analysis, following the manufacturer’s instructions. In brief, 1 × 10^7^ MDA-MB-231 cells were treated with 50 ng/ml IL-6 for 30 minutes and then cross-linked with 1% formaldehyde (Sigma-Aldrich). After fixation and lysis, chromatin was ultrasonically sheared under pre-optimized conditions and subsequently immunoprecipitated with Phospho-Stat3 antibody (Cell Signaling) or goat anti-human IgG at 4°C overnight. DNA fragments were extracted and purified from the pull-down complexes, and PCR was performed using a thermocycler in a total volume of 20 μL. The program included the following steps: 94°C for 2 minutes; 30 cycles of denaturation at 94°C for 20 seconds, annealing at 59°C for 30 seconds, and extension at 72°C for 30 seconds. Finally, PCR products were electrophoresed on a 2% agarose gel. The primer sequences are as follows: Forward primer-1: 5’ AGAAGTTCAGCGCGGGATAA 3’; Reverse primer-1: 5’ TCGTGGATTCTGTGACTTCCTC 3’; Forward primer-2: 5’ CATATGGGTCTGCTGCTGACTT 3’; Reverse primer-2: 5’ TACCTTCAGAGGGGTAAGAGC 3’.

### Dual-luciferase assay

The human PDL-1 promoter was cloned into the pGL3 luciferase reporter vector (Promega, Madison, WI, USA). 500 ng pGL3 luciferase reporter vector with wild type or mutated human PDL-1 promoter, along with the pRL-TK Renilla luciferase vector control (1:10 ratio) were co-transfected into 293Tn or MDA-MB-231 cells using 1.5 µl of Lipofectamine LTX System (Thermo Fisher Scientific). After transfection for 48 hours, 293Tn or MDA-MB-231 cells were treated with IL-6 for 4 hours, and then the luciferase activity was measured using a dual luciferase reporter assay according to the manufacturer’s instructions (Promega). Additionally, 500 ng pGL3 luciferase reporter vector with wild-type or mutated human PD-L1 promoter, along with the pRL-TK Renilla luciferase vector control, were co-transfected into 293Tn or MDA-MB-231 cells with 500 ng of Stat3 pcDNA3 vector using 1.5 µl of Lipofectamine LTX (Thermo Fisher Scientific). After co-transfection for 48 hours, the luciferase activity was measured using a dual luciferase reporter assay. The Renilla luciferase was used as an internal control, and the results were calculated for relative luciferase activity using the Firefly Luc/Renilla Luc ratio.

### Statistical analysis

GraphPad Prism version 7.0 were utilized for data analysis. Tumor volume were exhibited with the mean plus and minus standard error of the mean, and one-way ANOVA were used to analyze the difference of tumor size at the endpoint. Student t test was used to evaluate the difference of tumor weight, percentage of Granzyme B+/CD8+ T cells and percentage of organoid viability.

## Results

### Sulindac improves the efficacy of anti–PD-L1 therapy *in vivo*

A syngeneic mouse model was established through mammary fat pad injection of 4T1 cells. Mice in each group were treated with either vehicle control CMC plus IgG2b, sulindac alone (7.5 mg/kg, p.o., b.i.d.), anti-mouse PD-L1 antibody alone (50 μg, i.p. b.i.w.), or a combination of sulindac and anti-mouse PD-L1 antibody (7.5 mg/kg p.o., b.i.d. + 50 μg, i.p. b.i.w.), respectively. Tumor progression was measured using an external caliper twice per week. As shown in Figure 1a, sulindac monotherapy and anti-mouse PD-L1 antibody monotherapy significantly reduced tumor growth in mice compared to vehicle control. However, the combination of sulindac and anti-mouse PD-L1 antibody further decreased tumor progression, which was significantly different from the sulindac or anti-mouse PD-L1 antibody monotherapy. Mice were sacrificed on day 27, and tumors were fully resected, removed, and weighed. Figure 1b and Supplementary Figure 1 show the average weight of 4T1 tumors at the endpoints. The results indicated that mice treated with the combination of sulindac and anti-mouse PD-L1 antibody had the smallest tumors compared to vehicle controls and sulindac or anti-mouse PD-L1 antibody monotherapy (p<0.05, t-test), which is consistent with the results of 4T1 tumor progression. Total body weight of mice was also measured twice per week, but no significant differences were observed between groups (Supplementary Figure 4a), supporting the safety of our treatment regimens in the 4T1 syngeneic mouse model. We also employed a second PBMC humanized mouse model to further confirm our results. The PBMC humanized mouse model was established by intraperitoneally injecting human peripheral blood mononuclear cells (PBMCs), isolated from healthy adult donors. Two weeks after PBMCs injection, we collected 40 μl of blood from the submandibular vein of mice in each group and measured the ratio of human CD45 immune cells and CD8 T cells in the blood using flow cytometry. The results in Supplementary Figure 2a and 2b showed that there were more than 60% human CD45+ immune cells and over 10% CD8+ T cells in the blood of PBMC humanized mice, indicating successful engraftment of human immune cells, including CD8+ T cells, in this mouse model. We treated each group of mice with either vehicle control CMC, sulindac alone (7.5 mg/kg, p.o., b.i.d.), anti-human PD-L1 antibody alone (200 μg, i.v. q.week), or a combination of sulindac and anti-human PD-L1 antibody (7.5 mg/kg p.o., b.i.d. + 200 μg, i.v. q.week), respectively. As shown in Figure 1c, compared with vehicle control or sulindac or anti-PD-L1 monotherapy, the combination of sulindac and anti-human PD-L1 antibody significantly reduced 56S PDX growth (p<0.05, t-test). We sacrificed mice four weeks after human PBMC injection, and tumors were collected and weighed at the endpoints. The results from two repeated PBMC humanized mouse experiments (Figure 1d and Supplementary Figures 3a and 3b) indicated that the combination of sulindac and anti-human PD-L1 antibody significantly reduced 56S tumor weight compared to vehicle controls and sulindac or anti-human PD-L1 antibody monotherapy (p<0.05, t-test), consistent with the results of 56S tumor growth. Total body weight measurements of mice revealed no significant differences between groups (Supplementary Figure 4b), supporting the safety of our treatment regimens in the PBMC humanized mouse model.

**Figure 1.**
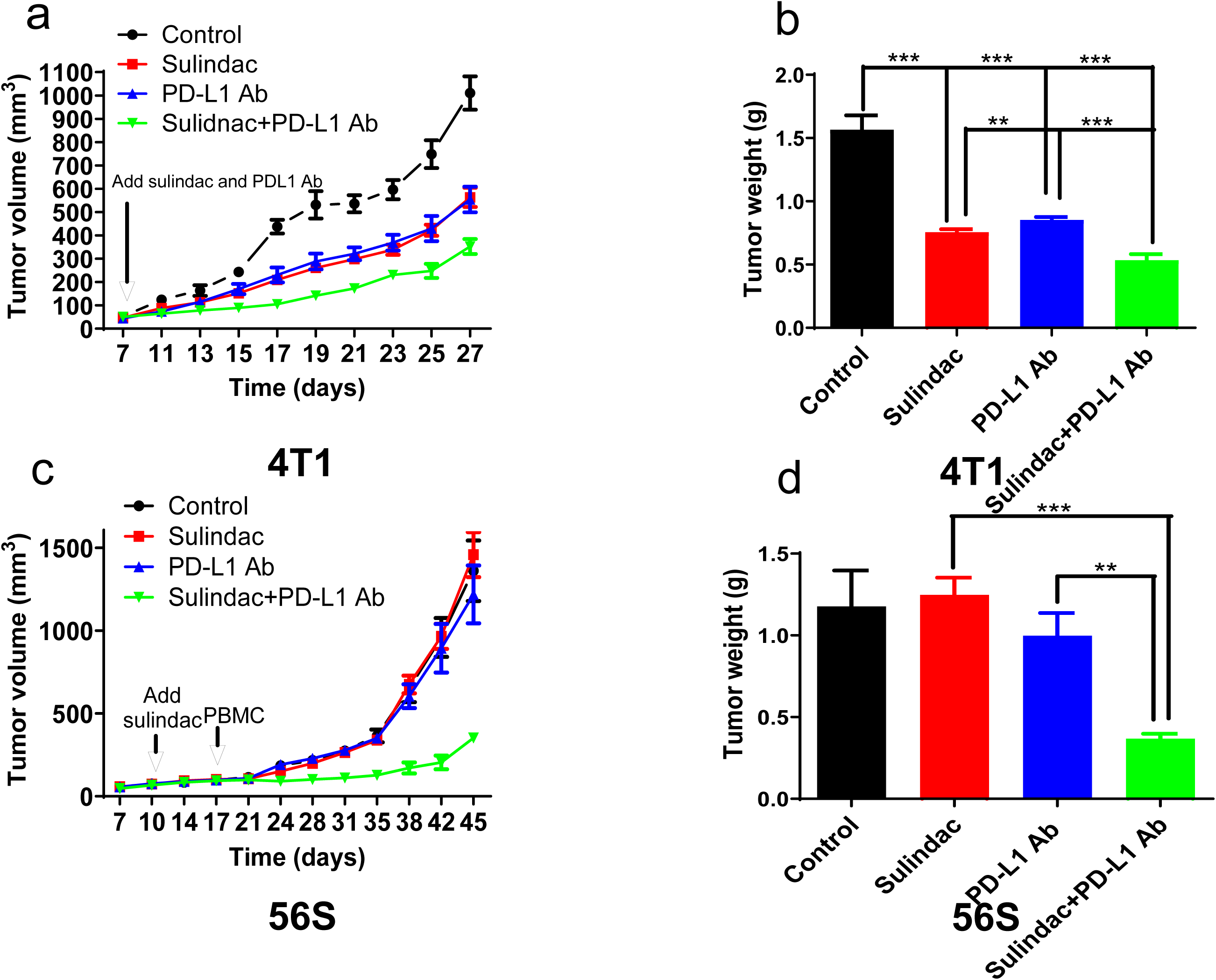
The combination of sulindac and PD-L1 antibody significantly inhibits breast tumor growth. (a) Monitoring 4T1 tumor orthotopic growth with external caliper measurements. Mice were screened with external caliper measurements twice per week after mammary fat pad injection. When orthotopic tumors reach about 50 mm^3^, twenty-eight female BALB/c mice were randomly divided into four groups: (1) vehicle control treated with CMC + IgG2b; (2) sulindac (7.5 mg/kg, p.o., bid.); (3) anti-mouse PD-L1 antibody (50 μg, i.p. b.i.w.); (4) combination with sulindac and PD-L1 antibody (7.5 mg/kg p.o., bid. + 50 μg, i.p. b.i.w.). BALB/c mice were sacrificed on day 27 after 4T1 cells injection. (b) 4T1 tumor weights of all mice at the endpoint. Differences between two groups were assessed by Student’s t-test, *p<0.05. (c) Monitoring 56S tumor orthotopic growth with external caliper measurements. Mice were screened with external caliper measurements twice per week after implantation of tumor tissues into mammary fat pad. When orthotopic tumors reach about 70 mm^3^, mice were pretreated with sulindac for up to 1 week, and 2×10^7^ human PBMCs were intraperitoneally injected into each mouse. Then twenty-four female NOD/SCID mice were randomly divided into four groups: (1) vehicle control treated with CMC; (2) sulindac (7.5 mg/kg, p.o., b.i.d.); (3) anti-human PD-L1 antibody (200 μg, i.v. q.week); (4) combination with sulindac and anti-human PD-L1 antibody (7.5 mg/kg p.o., b.i.d. + 200 μg, i.v. q.week). NOD/SCID mice were sacrificed after human PBMCs injection for up to 4 weeks. (d) 56S tumor weights of all mice at the endpoint. Differences between two groups were assessed by Student’s t-test, *p<0.05.

### Sulindac increases the infiltration of CD8+ T lymphocytes in tumor tissues by downregulating the expression of the PD-L1 gene

A previous study demonstrated that CD8+ T lymphocytes play a crucial role in sulindac’s inhibitory activity on tumor cell growth, particularly in a 4T1 syngeneic mouse model [22]. We collected 4T1 tumor tissues to analyze mouse CD8+ T lymphocyte infiltration using flow cytometry and immunofluorescence staining. As shown in Figure 2a, sulindac and PD-L1 antibody treatment significantly increased activated mouse CD8+ T lymphocytes (Granzyme B+/CD8+) in the tumor tissues. The combination group treated with sulindac and PD-L1 antibody exhibited the highest infiltration of activated mouse CD8+ T lymphocytes in the tumor tissues. Immunofluorescence staining (Figure 3a and supplementary Figure 5) of 4T1 tumor also suggests that the combination of sulindac and PD-L1 antibody significantly increased the infiltrating mouse CD8+ T lymphocytes in the 4T1 tumor tissues. These results support the idea that sulindac can promote active mouse CD8+ T lymphocyte infiltration in mouse breast tumors. We also collected human 56S PDX tumor tissues and tumor organoids to analyze human CD8+ T lymphocyte infiltration using flow cytometry and immunofluorescence staining. As shown in Figure 2b, sulindac treatment slightly increased activated human CD8+ T lymphocytes (Granzyme B+/CD8+) in the tumor tissues, while the combination of sulindac and PD-L1 antibody significantly increased the infiltrating active human CD8+ T lymphocytes in the 56S tumor tissues. Immunofluorescence staining (Figure 3b and 4a; Supplementary Figure 6 and 7) of 56S tumor tissues and organoids revealed that 56S tumor tissues or organoids treated with sulindac had increased human CD8+ T lymphocyte infiltration, and organoids treated with a combination of sulindac and anti-human PD-L1 antibody had the highest infiltration of human CD8+ T lymphocytes. Furthermore, we treated the 56S control tumor organoids, mixed with 1×10^6^ activated human PBMC, with a low dose of SS (sulindac sulfide, 20µM) and 200 nM anti-human PD-L1 antibody for 48 hours, after which we detected CD8+ T lymphocyte staining in the 56S tumor organoids. As shown in Figure 4b and Supplementary Figure 8, SS and the combination of SS with anti-human PD-L1 antibody increased human CD8+ T lymphocytes in the 56S tumor organoids, further supporting that sulindac can enhance human CD8+ T lymphocyte infiltration in human 56S tumors. Additionally, we employed a tumor organoid cell viability assay to investigate whether sulindac could affect tumor organoid cell viability. We treated 56S tumor organoids (with or without activated human PBMC) with 0.1% vehicle control DMSO, 20 µM sulindac sulfide (SS), 200 nM anti-human PD-L1 antibody, and a combination of 20 µM SS with 200 nM anti-human PD-L1 antibody. The results of *in vitro* organoid cell viability assay (Figure 5a and 5b) indicated that the combination of sulindac with PD-L1 antibody, along with activated human PBMC, significantly reduced 56S organoid cell viability but did not affect 56S tumor organoid viability in the absence of activated PBMC. This supports the notion that sulindac can decrease 56S organoid cell viability by facilitating the infiltration of human CD8+ T lymphocytes. Consistent with the results of our previous publication [23], when analyzing PD-L1 expression in the same set of tumor samples, we found that PD-L1 expression was decrease when infiltrating CD8+ T lymphocytes in all groups (Figure 6a and 6b; Supplementary Figure 9 and 10).

**Figure 2.**
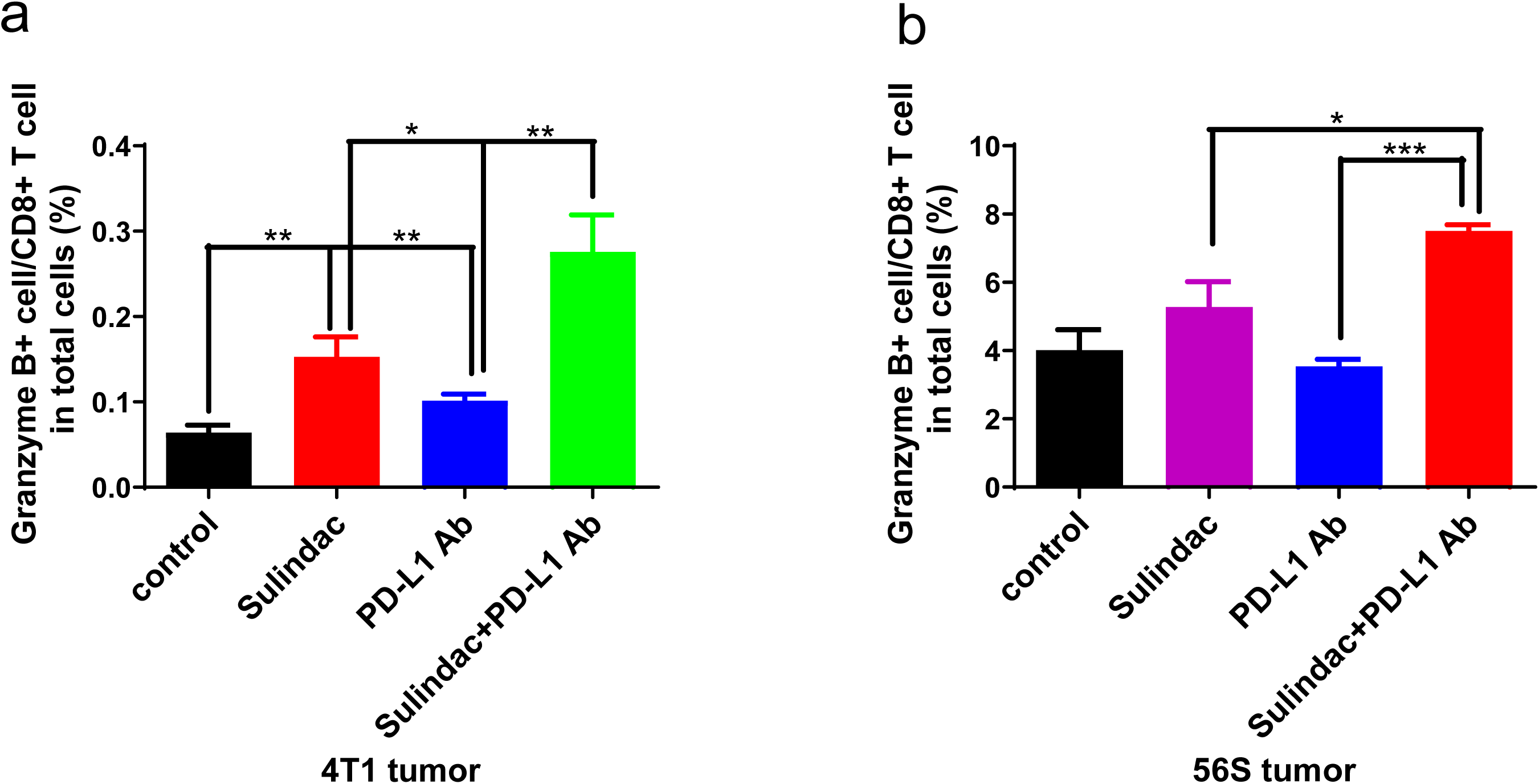
The combination of sulindac and PD-L1 antibody significantly increase infiltration of activated CD8+ T lymphocytes in the tumor tissues. We prepared single-cell suspensions from the 4T1 and 56S tumor through mechanical dissociation as described in Materials and Methods. Then cells were incubated with flow cytometry antibodies as described in Materials and Methods prior to the analysis with flow cytometry.

**Figure 3.**
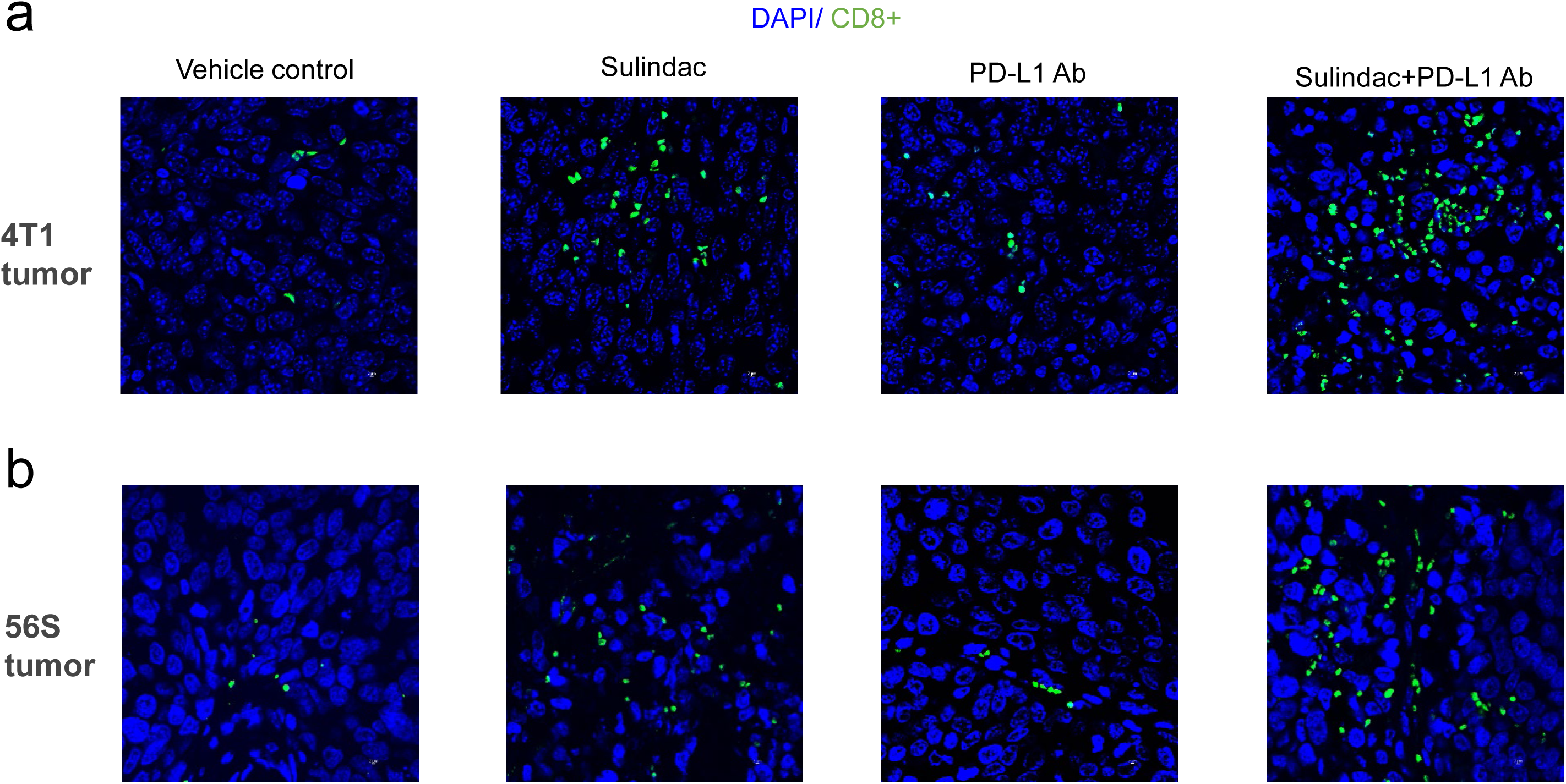
Immunofluorescence imaging results show that sulindac increases the infiltration of CD8+ T lymphocytes in 4T1 and 56S tumor tissues. Treatments include vehicle control, sulindac, PD-L1 Ab, and sulindac+PD-L1 Ab. Green: CD8+ T cells; Blue: DAPI.

**Figure 4.**
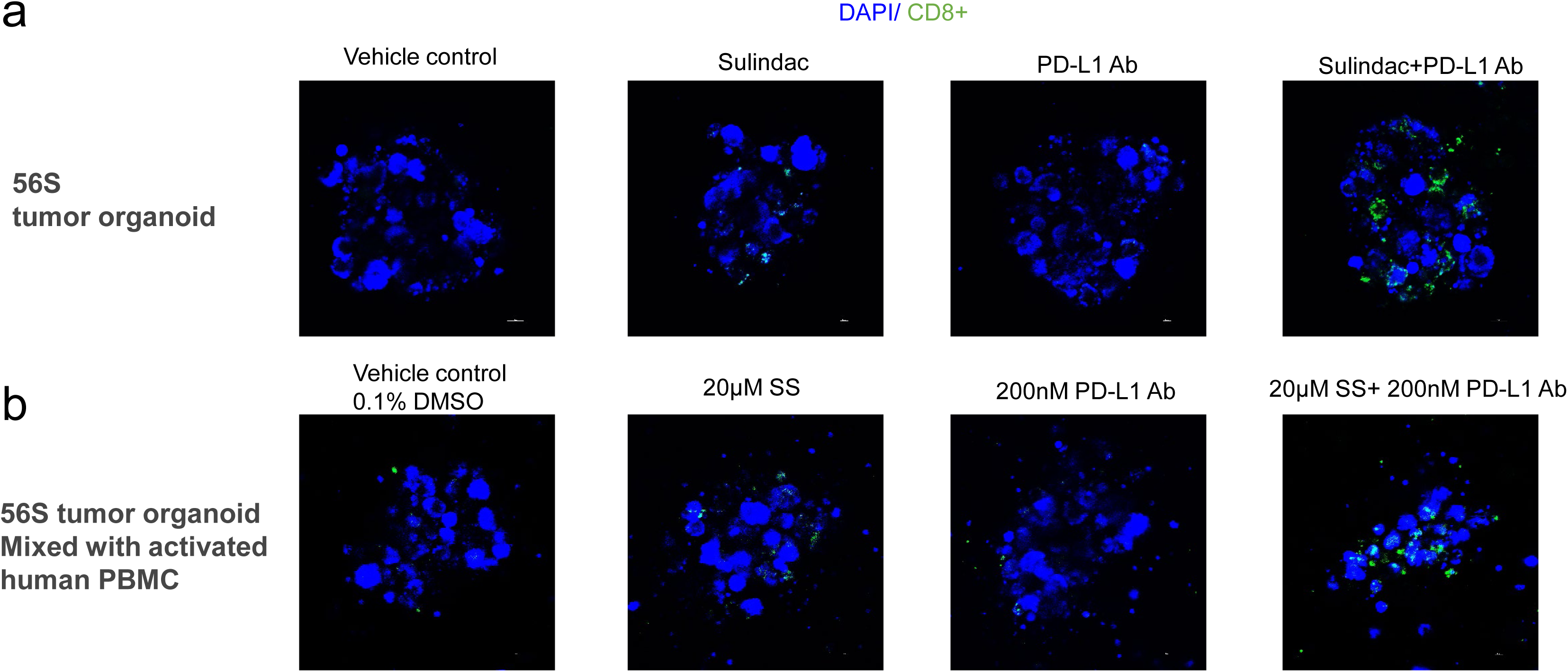
Immunofluorescence imaging results show that sulindac increases the infiltration of CD8+ T lymphocytes in 56S organoid. (a) After isolation of the 56S tumor organoid, the organoids were incubated with FITC mouse anti-human CD8 antibody for overnight at 37°C and 5% CO2 in a humidified incubator, followed by 5 µl/ml of Hoechst 33342 solution for 30 min. Treatments include vehicle control, sulindac, PD-L1 Ab, and sulindac+PD-L1 Ab. Green: CD8+ T cells; Blue: Hoechst 33342. (b) After the 56S control tumor organoids were mixed with 1×10^6^ activated human PBMCs, the tumor organoids were treated with vehicle control DMSO at 0.1%, 20 µM sulindac sulfide (SS), 200 nM anti-human PD-L1 antibody (Atezolizumab), and combination of 20 µM SS with 200 nM anti-human PD-L1 antibody (Atezolizumab) for 2 days. The 56S tumor organoids were then incubated with FITC mouse anti-human CD8 for overnight at 37°C and 5% CO_2_ in a humidified incubator, followed by 5 µ/ml of Hoechst 33342 solution for 30 min

**Figure 5.**
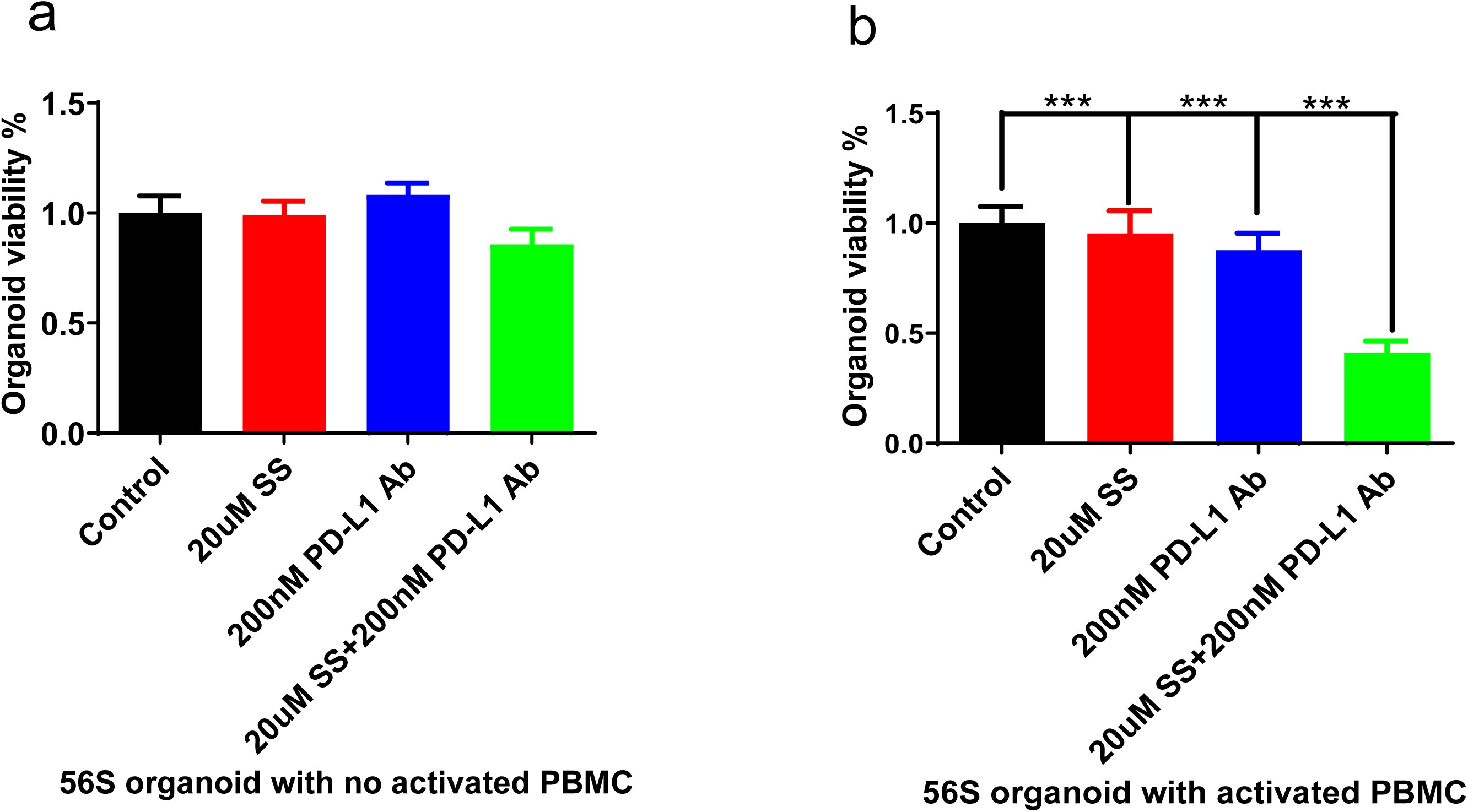
Tumor organoid cell viability assay results show that the combination of sulindac with PD-L1 antibody, along with activated human PBMC, significantly reduced 56S organoid cell viability. Referring to a published protocol [20], we treated 56S tumor organoid (with addition of either non-activated or activated human PBMCs.) with 0.1% DMSO vehicle control, 20 µM sulindac sulfide (SS), 200 nM anti-human PD-L1 antibody (Atezolizumab), and a combination of 20 µM SS with 200 nM anti-human PD-L1 antibody (Atezolizumab) and incubated for an additional 72 hours. Then we analyzed tumor organoid viability by using Cell Titer-Glo Assay.

**Figure 6.**
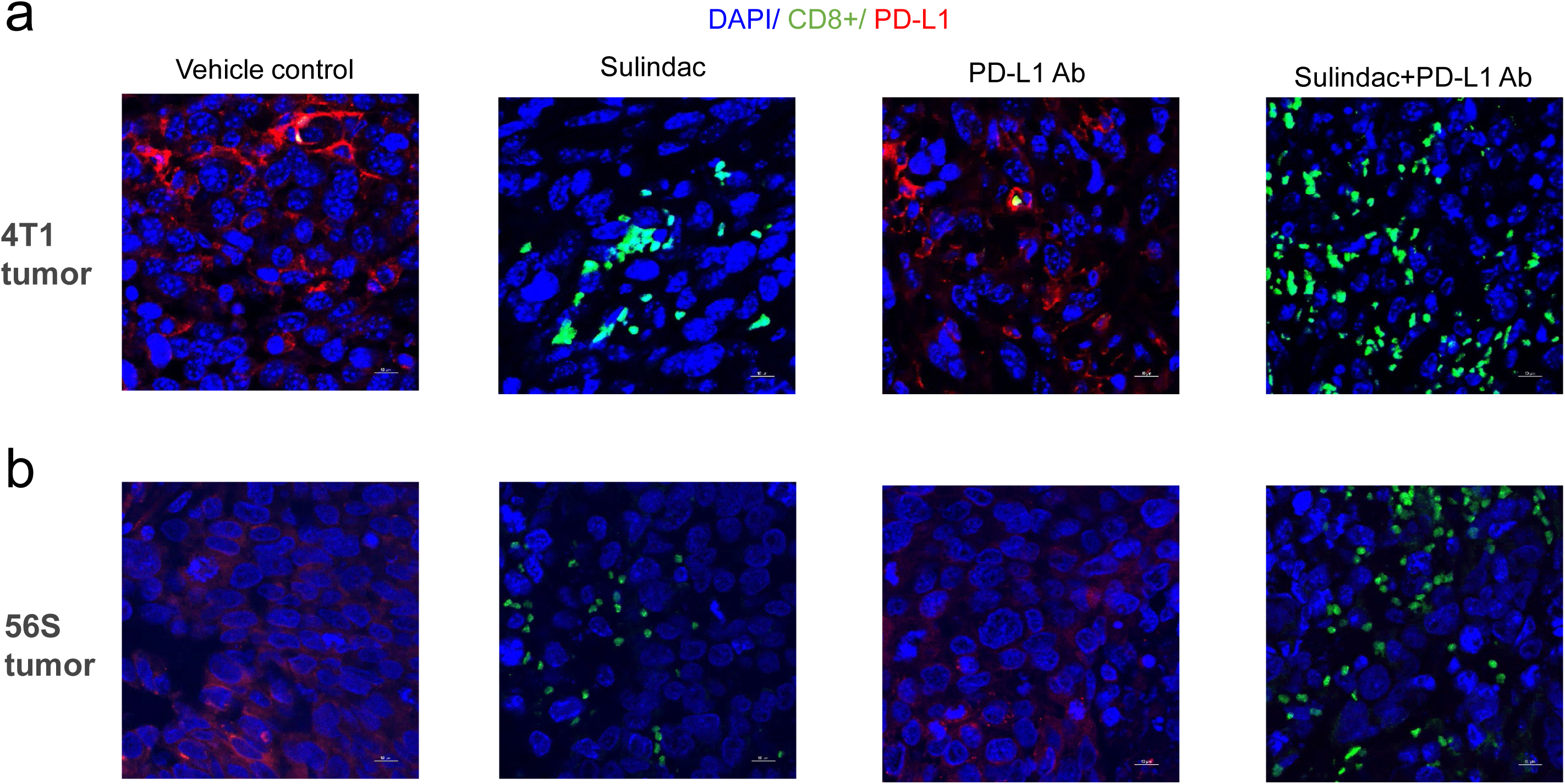
Sulindac increases the infiltration of CD8+ T lymphocytes in breast tumor tissues by down-regulating PD-L1 expression. Immunofluorescence imaging results show that sulindac increases the infiltration of CD8+ T lymphocytes in breast tumor tissues and downregulates PD-L1 expression. Treatments include vehicle control, sulindac, PD-L1 Ab, and sulindac+PD-L1 Ab. Red: PD-L1; Green: CD8+ T cells; Blue: DAPI. Images were taken with a Nikon Eclipse Ti2 Laser Confocal Scanning Microscope.

### Sulindac inhibits exosomal PD-L1 by downregulating the expression of the nsMase2 gene

Previous studies have demonstrated that circulating exosomal PD-L1 can bind to and consume PD-L1 antibodies [24, 25]. After extracting exosomes from mouse plasma samples of the 4T1 syngeneic mouse model and the PBMC humanized mouse model, we examined mouse and human exosomal PD-L1 expression in each treatment group. We also measured exosome particle concentration using NanoSight NS300. The number of exosome particles and exosome markers CD63 and CD9 were employed as endogenous controls to normalize the expression of mouse and human exosomal PD-L1, respectively. As shown in Figure 7a and 7b, the results from both mouse models were consistent. The expression of mouse or human exosomal PD-L1 was reduced in all treated mice compared to the vehicle control group, with the combination group displaying the lowest mouse or human exosomal PD-L1 expression. Supplementary Figures 11 and 12 show the results of Western Blot and relative quantitation of mouse or human exosomal PD-L1 in mice. To elucidate the mechanism of exosomal PD-L1 inhibition by sulindac, we focused on nsMase2, which plays a major role in the production of exosomal PD-L1 [25, 26]. We established nsMase2 knockdown breast cancer cell lines using CRISP/Cas9 technology and examined intracellular PD-L1 and exosomal PD-L1 expression following SS treatment. As shown in Figures 8a, 8b, 9a, 9b and 10, the results indicated that SS treatment, consistent with nsMase2 knockdown as a positive control, could decrease intracellular PD-L1 and exosomal PD-L1 expression without affecting exosome particle concentration in the human breast cancer cell lines, MDA-MB-231 and HCC70. These findings suggest that sulindac inhibits exosomal PD-L1 by downregulating the expression of the nsMase2 gene.

**Figure 7.**
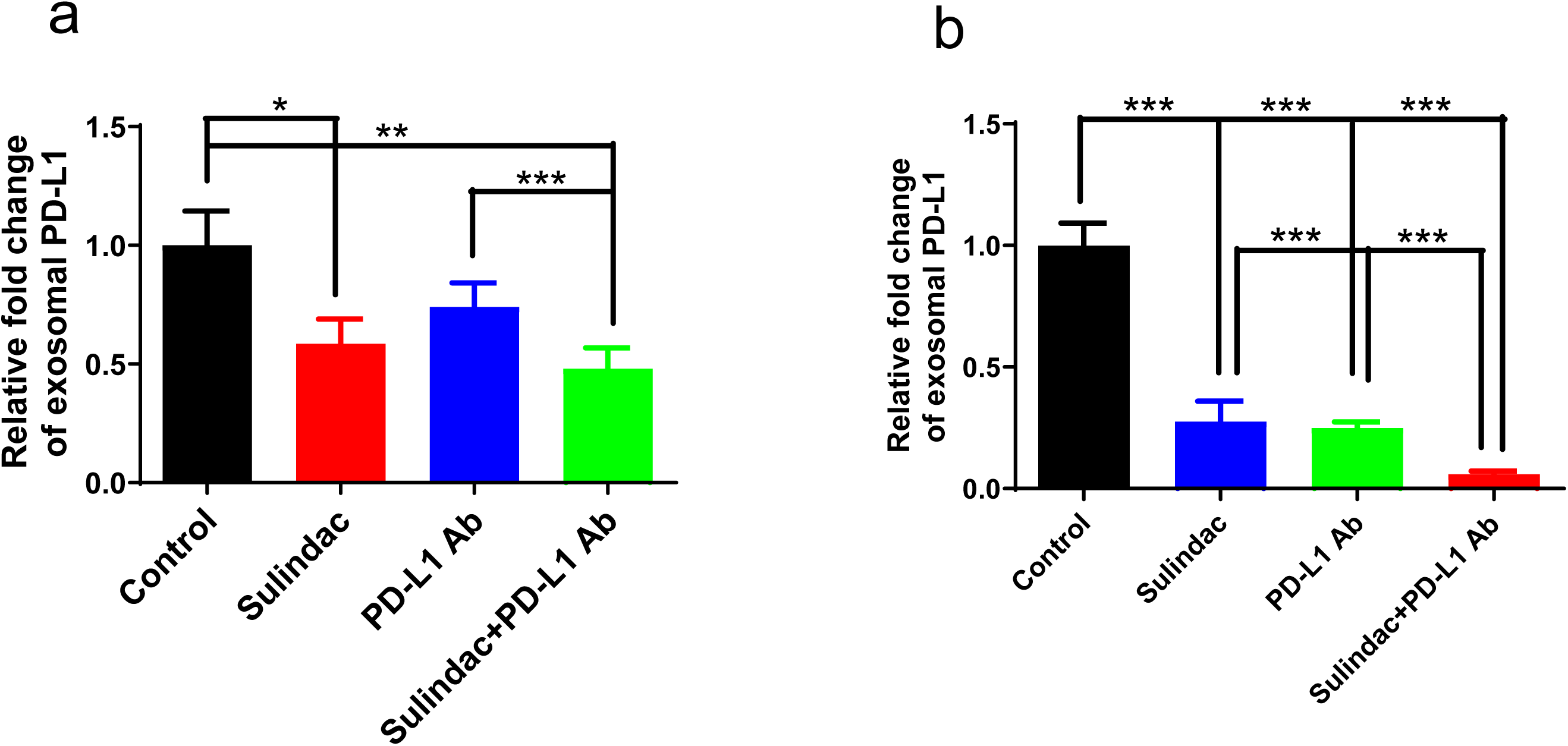
Sulindac inhibits exosomal PD-L1. (a-b) Expression of exosomal PD-L1 was decreased in mice treated with all drugs compared to vehicle controls, and the combination of sulindac and PD-L1 Ab showed the lowest expression of exosomal PD-L1. Exosomes were extracted from mouse plasma samples, purified with ExoQuick® ULTRA EV isolation kit, and lysed using RIPA buffer. Mouse exosomal PD-L1, human exosomal PD-L1, CD63 and CD9 were detected by Western blotting. The number of exosome particles and exosome markers CD63 and CD9 were employed as endogenous controls to normalize the expression of mouse and human exosomal PD-L1. Quantity One software (Bio-Rad) was used to calculate the intensity ratio between mouse exosomal PD-L1 and CD63 or human exosomal PD-L1 and CD9.

**Figure 8.**
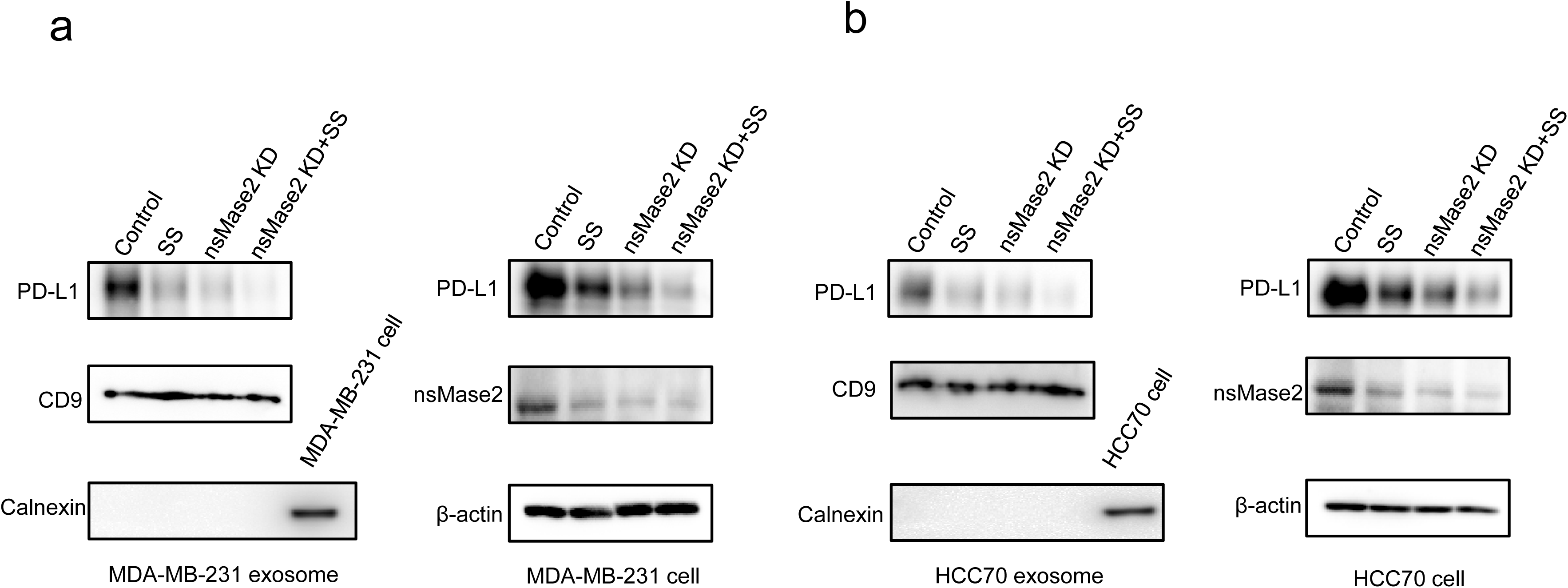
Sulindac inhibits exosomal PD-L1 by downregulating the expression of the nsMase2 gene. (a-b) After MDA-MB-231-control or HCC70-control cells and MDA-MB-231-nsMase2 KD or HCC70-nsMase2 KD cells were treated with 20uM SS for 48h, total cell lysates were prepared with RIPA buffer and exosome from cell culture media were isolated and purified with ExoQuick-TC PLUS Exosome Purification Kit. Then exosome lysates were generated using RIPA buffer. Lastly, the intracellular PD-L1, nsMasae2 and exosomal PD-L1 were analyzed by Western blot.

**Figure 9.**
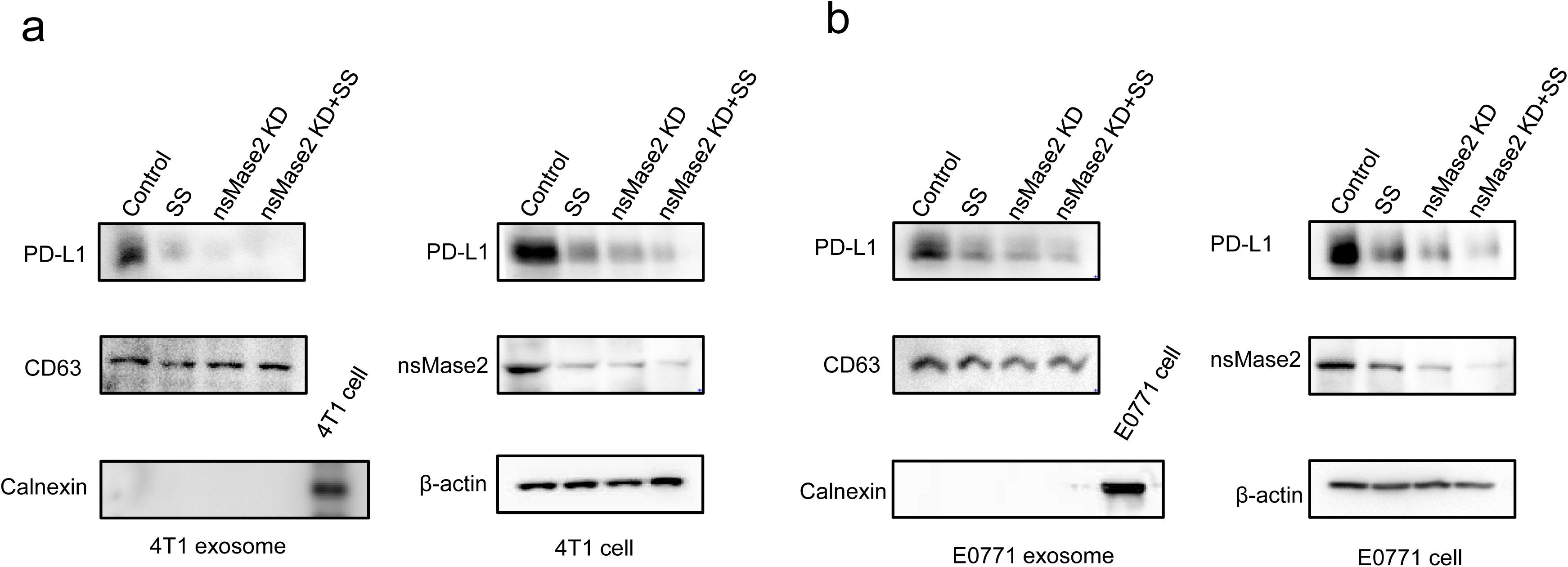
Sulindac inhibits exosomal PD-L1 by downregulating the expression of the nsMase2 gene. (a-b) After 4T1-control or E0771-control cells and 4T1-nsMase2 KD or E0771-nsMase2 KD cells were treated with 20uM SS for 48h, total cell lysates were prepared with RIPA buffer and exosome from cell culture media were isolated and purified with ExoQuick-TC PLUS Exosome Purification Kit. Then exosome lysates were generated using RIPA buffer. Lastly, the intracellular PD-L1, nsMasae2 and exosomal PD-L1 were analyzed by Western blot.

**Figure 10.**
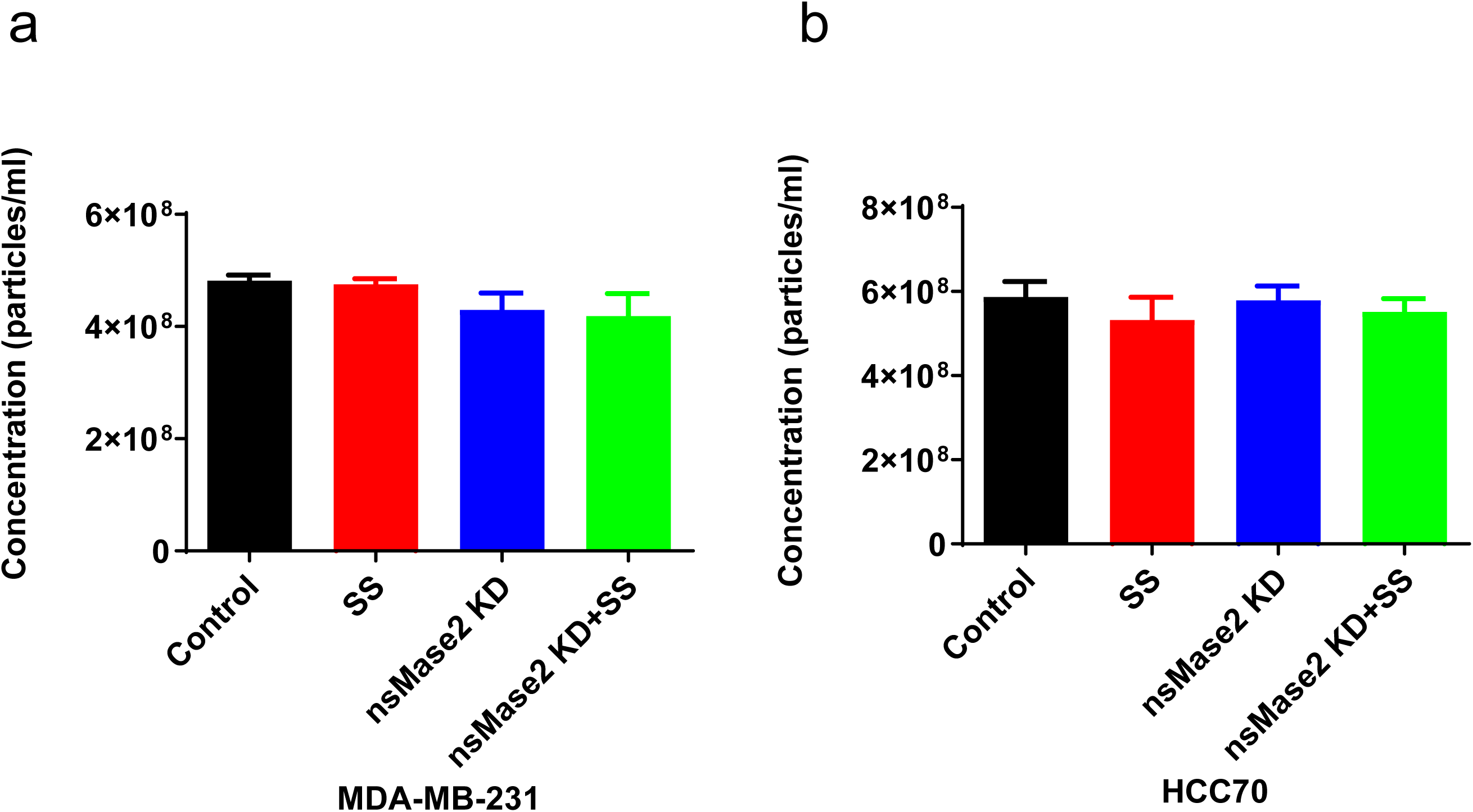
Sulindac sulfide treatment can not affect total exosome secretion. After exosome from cell culture media were isolated and purified with ExoQuick-TC PLUS Exosome Purification Kit, exosome concentrations were measured using the NanoSight NS300

### Blockade of Stat3 signaling is associated with sulindac modulation of TNBC cells response to anti-PD-L1 therapy

Previous studies have demonstrated that sulindac inhibits the Stat3 signaling pathway [27–29]. Recent studies also reported the ability of Stat3 to regulate PD-L1 in various cancer cells [30–32]. Since our *in vivo* data showed that sulindac could downregulate PD-L1 expression in tumor tissues, we investigated whether Stat3 signaling might be involved in sulindac’s inhibitory effect on PD-L1 expression in TNBC cells. Figure 11a presents two putative binding sites for Stat3 (GCTTCCGCAGC; AGGCTTTTATCAGAAAGGGGG) within the promoter region of the PD-L1 gene, as reported previously [33, 34]. We further determined the binding of Phospho-Stat3 to the promoter of PD-L1 using the chromatin immunoprecipitation (ChIP) assay. Briefly, Phospho-Stat3 antibodies were used to immunoprecipitate the sheared chromatin fractions extracted from IL-6-treated MDA-MB-231 cells. After purifying the DNA fragments from the pull-down complexes, we performed PCR analysis with the designated primers to detect the target DNA fragments containing the putative Stat3 binding sites. As shown in Figure 11a and 11b, we detected a band with the expected size using DNA templates extracted from the Phospho-Stat3 immunoprecipitated complex. Additionally, we employed a dual-luciferase reporter assay to confirm the ChIP results. Briefly, wild-type or mutated promoter fragments of the human PD-L1 gene were cloned into the pGL3 luciferase reporter vector, and the dual-reporter system used Firefly Luciferase as the promoter reporter and Renilla as an internal control for signal normalization. After transfecting these constructs into MDA-MB-231 and 293Tn cells, we measured the signals of Firefly Luc and Renilla Luc and calculated the relative light unit (RLU) ratio. As shown in Supplementary Figure 13 and 14, the RLU of the mutated constructs was much lower than that of the wild-type constructs upon IL-6 treatment or Stat3 pcDNA3 co-transfection (p<0.05). Together, these results provide strong evidence that Phospho-Stat3 directly binds to the promoter of PD-L1 in MDA-MB-231 and 293Tn cells. Furthermore, we utilized immunofluorescence imaging to analyze the regulation of PD-L1 expression following sulindac treatment. IL-6 and the Stat3 inhibitor (Stattic) were included to stimulate and block Stat3 signaling, respectively Jochen [35, 36]. All four TNBC cell lines were used in this study. As shown in Figure 12 and Supplementary Figures 15-18, IL-6 treatment promoted the translocation of Phospho-Stat3 into the nucleus, which in turn upregulated PD-L1 expression. Additionally, treatment with SS effectively attenuated the induction of IL-6 and blocked Stat3 signaling, which was comparable to that of the Stat3 inhibitor Stattic. We also extended our study by examining Phospho-Stat3 and PD-L1 in 4T1 and 56S tumor tissues that we analyzed above. As expected, mice treated with sulindac alone or in combination with PD-L1 Ab exhibited reduced Stat3 signaling in the nucleus along with PD-L1 downregulation (Figure 13 and Supplementary Figure 19-20), consistent with the *in vitro* results.

**Figure 11.**
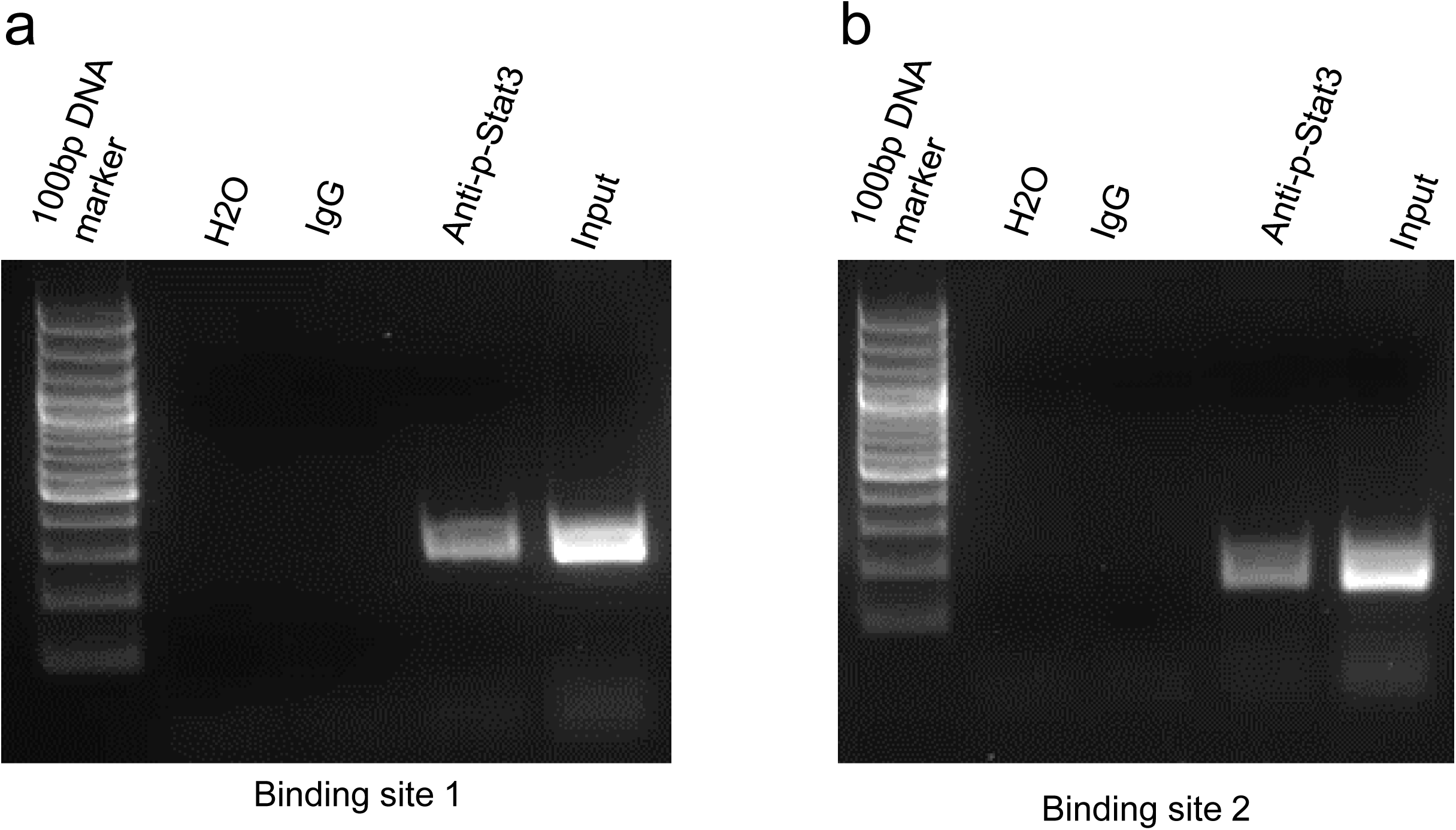
ChIP assay results showing direct binding of p-Stat3 to the PD-L1 promoter. MDA-MB-231 cells were pre-treated with 50 ng/ml IL6 for 30 min and immunoprecipitated by Phospho-Stat3 antibody or normal mouse IgG. The isolated DNA fragments were purified from the pull-down complexes and utilized as the templates for PCR amplification. The expected size of the PCR product containing the putative p-Stat3 binding sequence one and two is 267 bp and 191 bp, respectively. Input samples were derived from the starting chromatin that had been used for ChIP.

**Figure 12.**
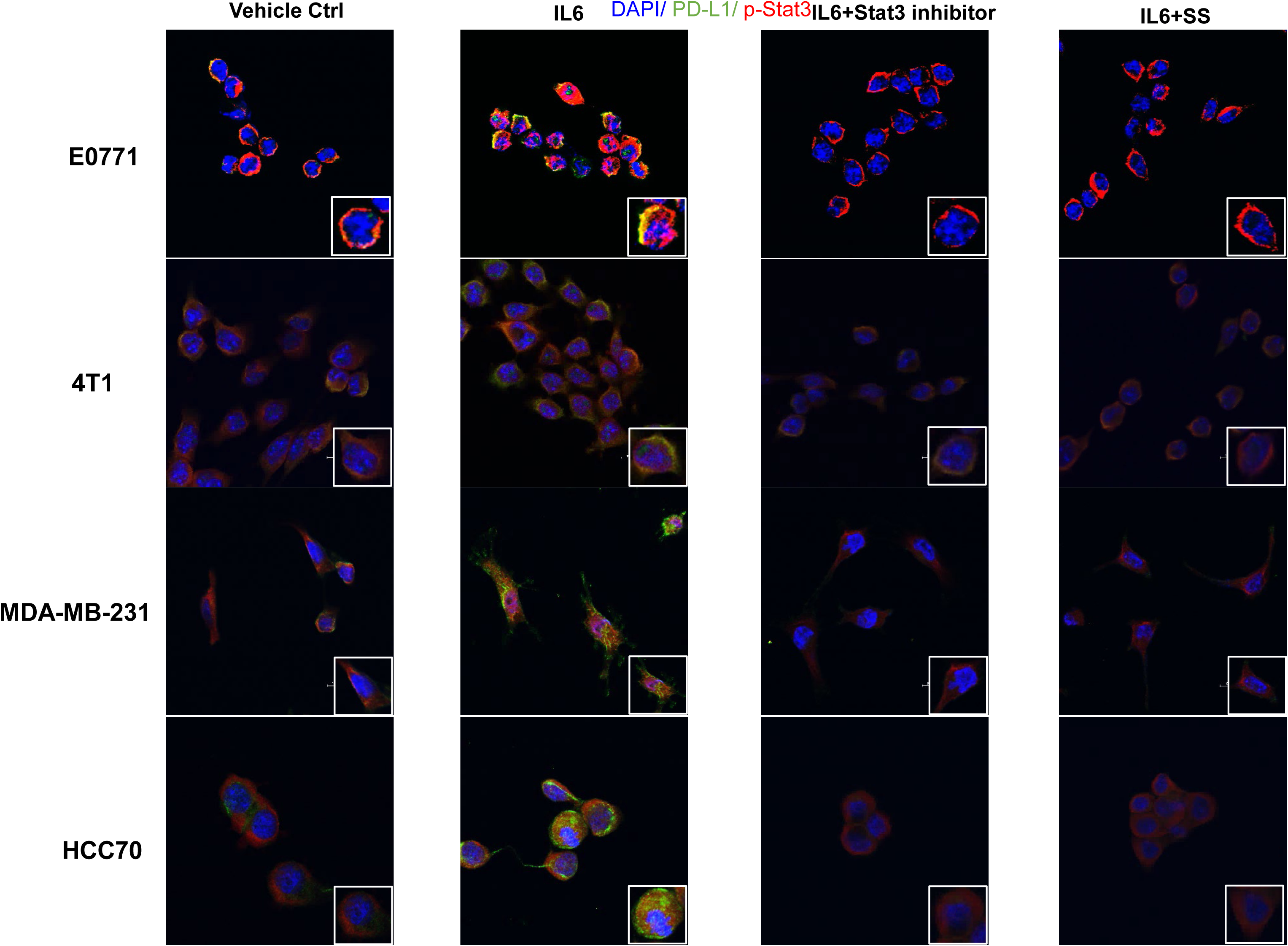
SS attenuates the induction of Stat3 signaling by IL6. We treated four triple negative breast cancer cell lines initially with a 10 μM concentration of a Stat3 inhibitor for two hours and a 20 μM dose of SS for twelve hours. As a vehicle control, DMSO was utilized. Then, either human or mouse IL6, at a concentration of 50 ng/ml, was introduced to these cells and incubated for thirty minutes. For fixation, the cells were treated with 4% formaldehyde for ten minutes and subsequently permeabilized using 1% Triton X-100. After a blocking phase with 1% BSA, the cells underwent overnight incubation at 4°C with PD-L1 and p-Stat3 antibodies. Following this, staining procedures were carried out, and the cells were visualized using confocal microscopy.

**Figure 13.**
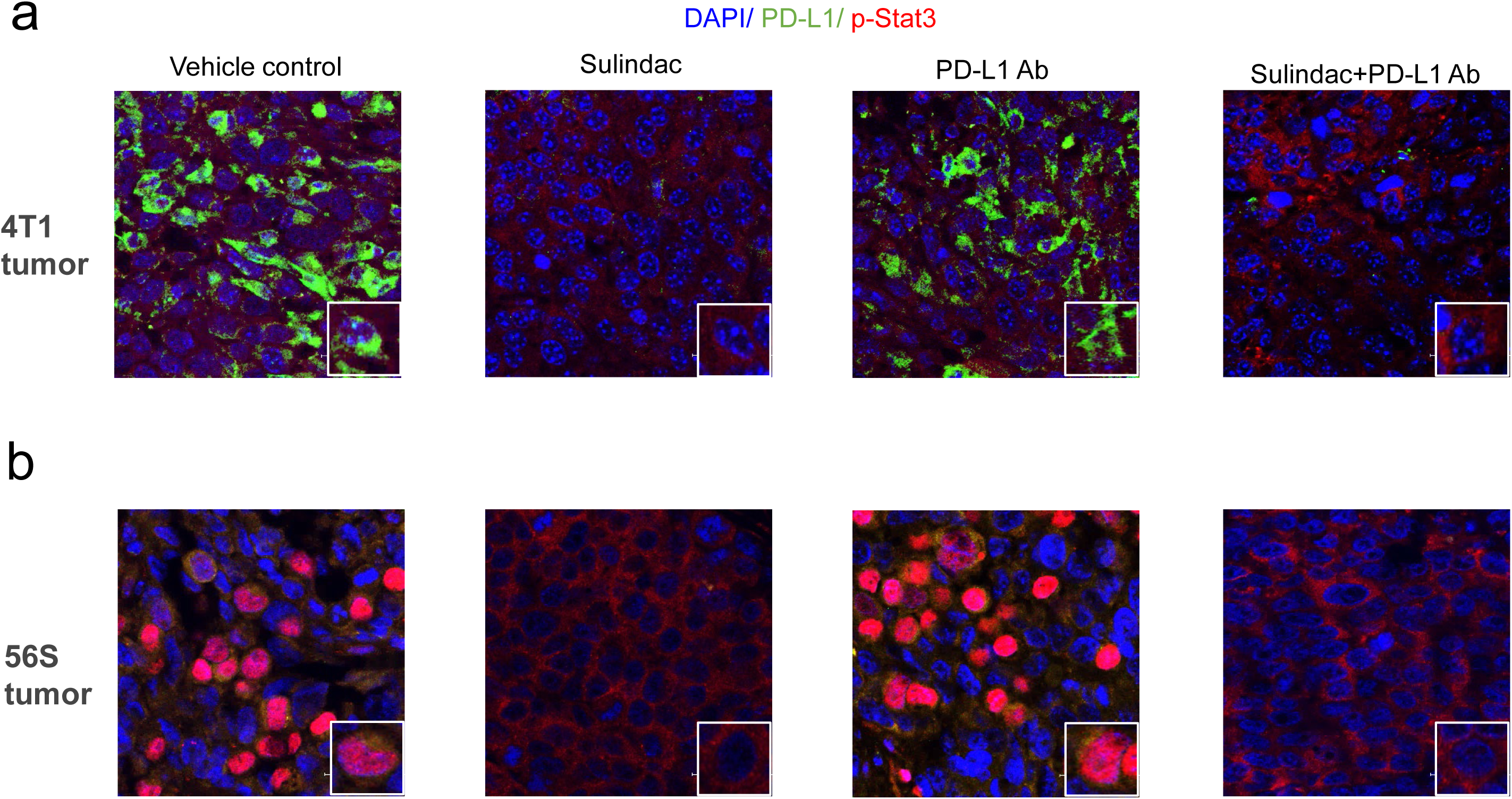
Sulindac downregulates PD-L1 expression by blocking Stat3 signaling in 4T1 and 56S tumor issues. (a-b) This analysis employed the same sample sets as those used in Figure 6. Red: p-Stat3; Green: PD-L1; Blue: DAPI. Images were captured by using a Nikon Eclipse Ti2 Laser Confocal Scanning Microscope.

### Hsa-miR-570 mediates sulindac sulfide inhibitory effect on PD-L1

Using miRNA prediction tools (Targetscan), hsa-miR-616, hsa-miR-570, and hsa-miR-548a were identified as potential miRNAs targeting PD-L1. We previously reported that SS upregulates these miRNAs in HCC116 [17]. In this study, we aimed to determine if these miRNAs are also involved in sulindac sulfide-mediated PD-L1 reduction. In HCC70 and BT549 cells after SS treatment, we examined the expression of hsa-miR-616, hsa-miR-570, and hsa-miR-548a, but found only hsa-miR-570 to be upregulated (Figure 14). One study also supports that has-miR-570 can target PD-L1 [37]. We therefore investigated whether SS inhibited PD-L1 through the upregulation of hsa-miR-570. As shown in Figure 15, overexpression of hsa-miR-570 by transfection of its mimics could decrease PD-L1 expression similarly to SS treatment. Additionally, the combination of hsa-miR-570 mimic transfection with SS treatment further decreased PD-L1 expression. Conversely, downregulation of hsa-miR-570 by transfection of its inhibitor increased PD-L1 expression. SS treatment attenuated the promoting effect of hsa-miR-570 inhibitor on PD-L1. These results suggest that hsa-miR-570 can mediate the inhibitory effect of SS on PD-L1.

**Figure 14.**
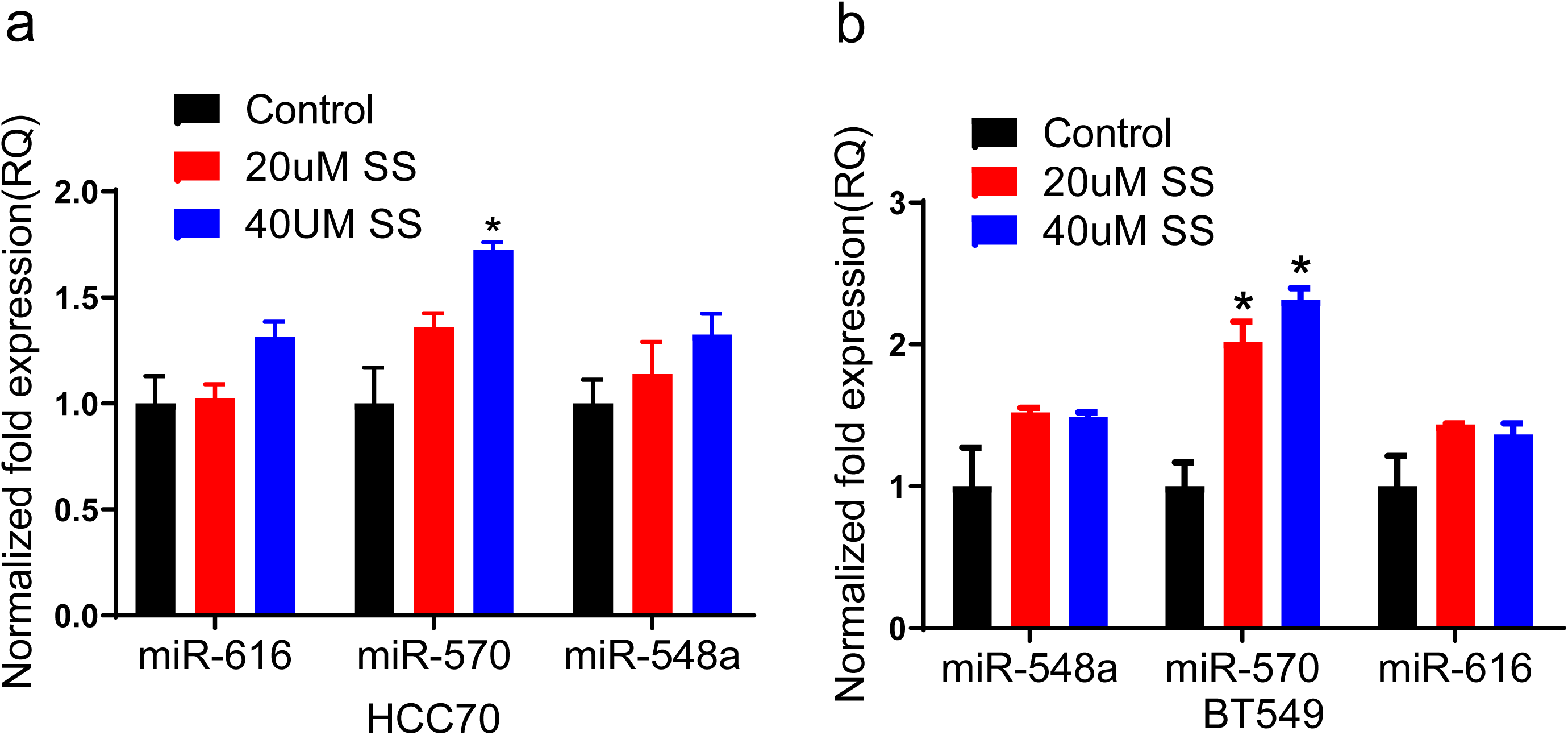
SS treatment significantly increase hsa-miR-570 in breast cancer cells HCC70 and BT549. After breast cancer cells were treated with SS for 48h, the expression of hsa-miR-616, hsa-miR-570 and hsa-miR-548 were examined by real-time PCR. * p<0.05.

**Figure 15.**
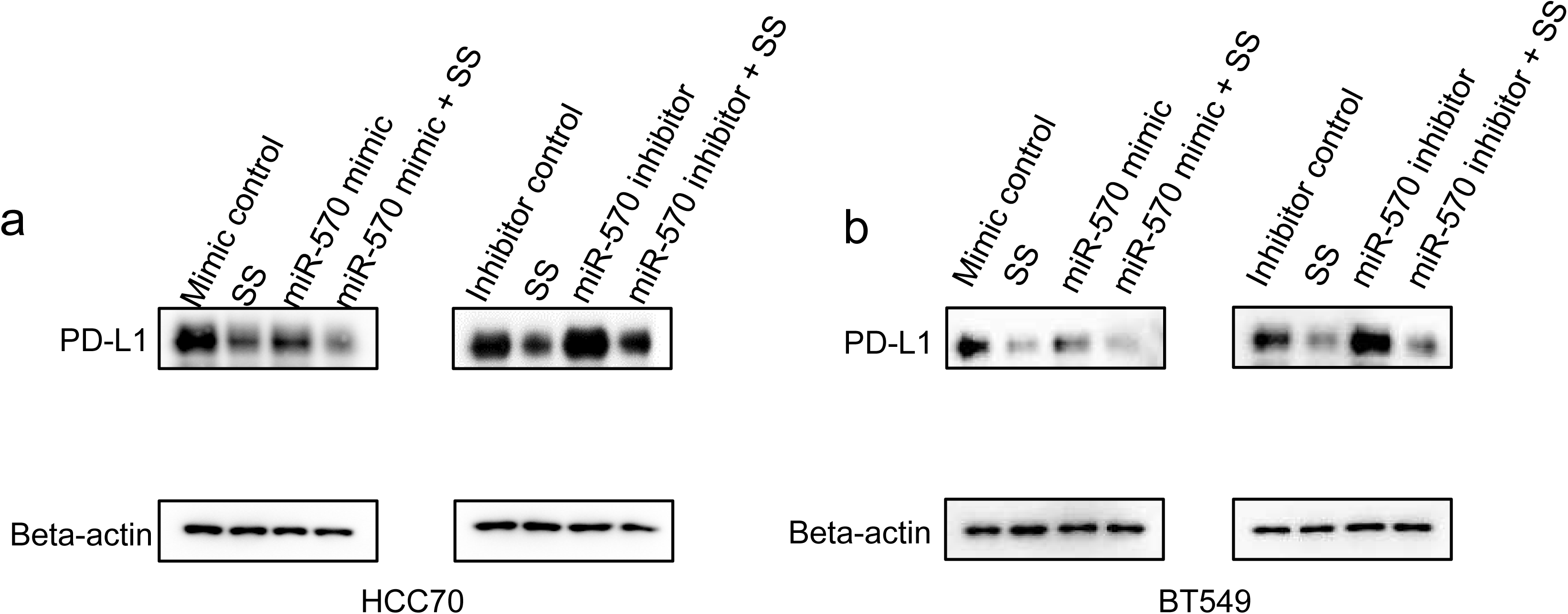
Hsa-miR-570 mediates sulindac sulfide inhibitory effect on PD-L1. After miR-570 mimic and inhibitor transfection for 24h, the breast cancer cells were treated with 20 μM SS for 48h and then the expression of PD-L1 was examined by Western-blot

### Low doses of sulindac can effectively inhibit PD-L1 without significant systematic toxicity

As mentioned in our previous publication [23], the FDA-approved dose of sulindac for human use is 300-400 mg/day. Referring to the published protocol [38], we calculated that equivalent dose for small animal studies ranges from 61.5-82.0 mg/kg. In this study, we treated mice with doses of only 7.5 mg/kg, which should be considered a low dose of sulindac and are equivalent to approximately 1/8 of the FDA-approved dose for human use. There have been concerns raised about possible human side effects of sulindac on the gastrointestinal, renal, and cardiovascular systems associated with COX-2/PGE2 inhibition [49]. Intriguingly, it has been reported that COX-2 is capable of impairing tumor immunity and that inhibition of COX-2 has the potential to improve immunotherapeutic responses in a variety of tumor models [50–52]. Therefore, sulindac holds great promise as an effective and safe addition to ICI therapy if the systemic toxicity directly associated with PGE2 inhibition can be minimized. To assess whether sulindac is systemically toxic, particularly regarding the inhibition of PGE2, we measured the concentration of PGE2 by ELISA using plasma samples collected from 4T1 and 56S mice. As shown in Figure 16a and 16b, there were no significant differences between the mice in each group in the two mouse models. We also monitored the total body weight of the mice (as a surrogate marker of toxicity) twice a week, we did not find any significant differences among the groups (Supplementary Figure 4). These results indicate that low-dose sulindac is a safe and effective option to sensitize TNBC to anti-PD-L1 immunotherapy.

**Figure 16.**
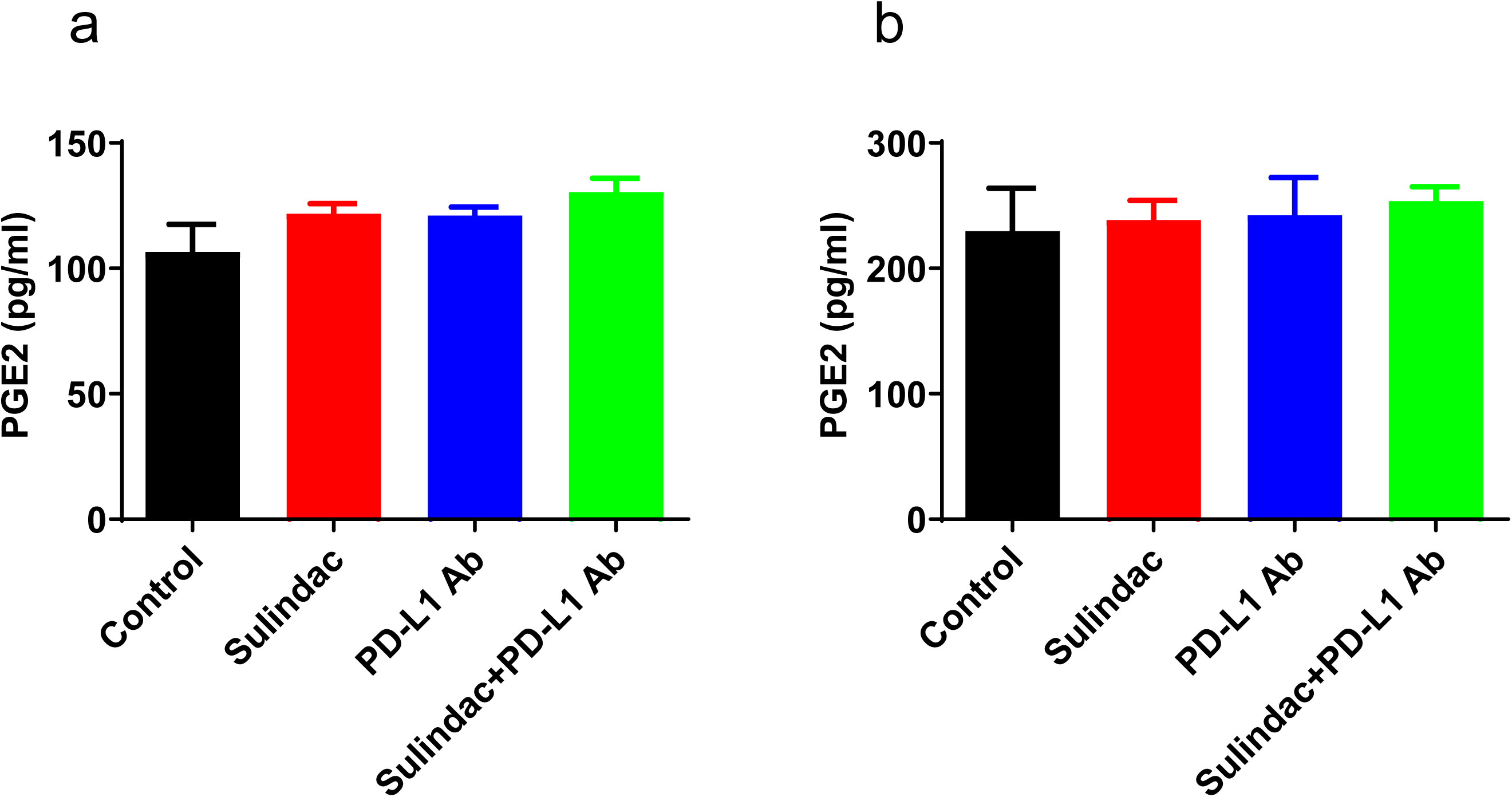
Low doses of sulindac have no significant impact on systemic PGE2. We measured PGE2 concentrations in 4T1 (a) and 56S (b) mouse plasma using an ELISA assay, employing the same plasma samples as those used in Figure 7. We diluted 50 μL of each plasma sample and placed them into a 96-well microplate coated with goat anti-mouse polyclonal antibodies. The samples were then incubated with 50 μL of a mouse PGE2 monoclonal antibody for one hour at ambient temperature. Subsequently, we added PGE2 conjugate to each well and allowed it to incubate for two hours with mild agitation. Following this, the wells were washed, and we added a substrate solution to each, allowing it to react for up to 30 minutes. To halt the color development, a stopping buffer was introduced. We then measured the absorbance at 450 nm using a Bio-Rad microplate reader.

## Discussion

The clinical efficacy of checkpoint immunotherapy has been recognized by experts in the field, particularly in treating metastatic melanoma and lung cancer, where it improves long-term and overall survival rates. Recently, its application has extended to metastatic triple-negative breast cancer [39]. However, clinical studies indicate that the benefits of ICI immunotherapy in treating TNBC are limited to a small fraction of patients with high PD-L1 levels, accounting for less than 38% of TNBC patients [40]. Unfortunately, for most TNBC patients, current ICI regimens offer minimal clinical benefit in terms of survival. Epidemiological studies have shown that nonsteroidal anti-inflammatory drugs (NSAIDs) significantly reduce breast cancer incidence and the risk of death[41, 42]. A recent study reported that breast cancer patients treated with the NSAID, ketorolac following mastectomy had up to 5-fold reduction of early relapse within a 9-to-18-month period compared with a non-treated group[43]. Our prior research also investigated the anticancer activity of sulindac and several derivatives in TNBC [17, 44, 45]. These studies support the feasibility by using sulindac to prevent disease recurrence and progression in TNBC patients.

We selected the murine TNBC cell line 4T1 and established a syngeneic animal model using immunocompetent mice. We also chose human TNBC tumor 56S and established a humanized animal model using immunodeficient mice by intraperitoneally injecting human peripheral blood mononuclear cells (PBMCs), isolated from healthy adult donors. After injecting 4T1 cells or 56S tumor tissue into the mammary fat pad of mice, we treated the mice with sulindac, PD-L1 antibody, or a combination of both. We monitored the orthotopic growth of 4T1 and 56S tumors by measuring tumor size. Our findings clearly demonstrate that the combination therapy of sulindac and PD-L1 antibody was the most effective, significantly impacting tumor weight, size, and regression.

Given a previous study reporting the involvement of CD8+ T lymphocytes in the anticancer activity of sulindac against breast cancer [22] and our previous findings showing sulindac increased CD8+ T lymphocyte infiltration in MSS colon tumors, we used flow cytometry and immunofluorescence imaging to detect the infiltration of CD8+ T lymphocytes, including activated Granzyme B+/CD8+ T lymphocytes, in TNBC tumors 4T1 and 56S. Our results indicated a significant increase in infiltrated activated Granzyme B+/CD8+ T lymphocytes in tumor tissues treated with sulindac alone and in combination, with the latter showing more intense T cell infiltration. Organoid cell viability assays further demonstrated that sulindac in combination with PD-L1 antibody and activated human PBMCs significantly reduced the cell viability of 56S organoids, supporting the notion that sulindac can reduce 56S organoid cell viability by promoting human CD8+ T lymphocyte infiltration.

We evaluated the expression of PD-L1 in four TNBC cell lines, E0771, 4T1, MDA-MB-231, and HCC70, after treatment with sulindac. Consistent with our previous findings in colon cancer, sulindac significantly downregulated PD-L1 levels in TNBC cells. We further explored the molecular mechanism of sulindac’s inhibition of PD-L1. Many reports suggest that sulindac can block Stat3 signaling [27–29], with Stat3 involved in the regulation of PD-L1 expression in cancer cells [30–32]. Following established protocols, we searched for Stat3 binding sequences and identified two putative sites in the PD-L1 gene promoter. Using ChIP and luciferase reporter assays, we confirmed direct binding between Stat3 and the PD-L1 promoter. Recent studies also found that IL6 can induce PD-L1 expression via the Stat3 pathway [36]. Thus, we used IL6 as an inducer to stimulate Stat3 signaling and then assessed whether sulindac treatment could effectively affect PD-L1 expression in the four TNBC cell lines. As expected, our results support the notion that Stat3 signaling is a key mechanism by which sulindac transcriptionally downregulates PD-L1. We further examined Stat3 translocation in 4T1 and 56S tumor tissues collected from *in vivo* studies. Consistent with *in vitro* cell results, treatment with sulindac alone or in combination with PD-L1 antibody significantly reduced nuclear Stat3 signaling intensity while downregulating PD-L1. We previously reported that SS upregulates certain tumor suppressor miRNAs in colon cancer cells. Using predictive tools, we identified tumor suppressor miRNAs, including hsa-miR-616, hsa-miR-570, and hsa-miR-548a, that could potentially target PD-L1. Therefore, we aimed to determine whether these miRNAs are also involved in SS-mediated PD-L1 reduction in TNBC cells. Our results support the notion that hsa-miR-570 is one of the mechanisms mediating SS’s inhibitory effect on PD-L1.

Exosomes are membrane vehicles ranging from 30 to 100 nanometers in size, released by many cell types during physiological processes. Importantly, exosomes facilitate intercellular communication by fusing with receptor cells and releasing their contents [46]. Tumor-derived exosomes Steinbichler [47] are often more abundant in cancer cells than in normal non-tumor cells and contain key oncogenic elements, including various miRNAs, mRNAs, proteins, and lipids, that can enter cells and influence signaling pathways related to tumor progression and metastasis Kahlert [48]. Recent studies reported that exosomal PD-L1 can bind to PD-L1 antibodies, thereby weakening the inhibitory effect of PD-L1 antibodies [24, 25]. We collected blood samples from two TNBC mouse models, isolated plasma samples, and analyzed exosomal PD-L1 in mice treated with sulindac (alone), PD-L1 Ab (alone), and in combination (sulindac plus PD-L1 Ab). Consistent with our previous publication, we found that all treatment regimens significantly reduced exosomal PD-L1 expression. To elucidate the mechanism of sulindac’s inhibition of exosomal PD-L1, we focused on nsMase2, which plays a major role in the generation of exosomal PD-L1 [25, 26]. We established nsMase2 knockout breast cancer cell lines and examined the expression of intracellular PD-L1 and exosomal PD-L1 following SS treatment. Our results indicate that, using nsMase2 gene knockout as a positive control, SS treatment can reduce the expression of both intracellular and exosomal PD-L1, but does not affect the concentration of exosomal particles in the two human breast cancer cell lines MDA-MB-231 and HCC70. These findings suggest that sulindac can inhibit the expression of exosomal PD-L1 by downregulating nsMase2 gene expression.

Sulindac, a nonsteroidal anti-inflammatory drug (NSAID) approved by the U.S. FDA, is a non-selective inhibitor of cyclooxygenase-1 and −2 (COX-1 and COX-2). Inhibition of COX-2 reduces the synthesis of prostaglandin E2 (PGE2), a key mediator of inflammation and immune regulation. Long-term use of NSAIDs, including high doses of sulindac, has been associated with reduced PGE2 levels, which can lead to adverse effects on the gastrointestinal tract, kidneys, and cardiovascular system [49]. However, COX-2 has also been implicated in tumor progression and immune evasion, with studies suggesting that its inhibition may enhance anti-tumor immune responses in various cancer models [50–52]. Therefore, if the systemic toxicity caused by inhibition of PGE2 can be minimized, sulindac could be a very safe and effective addition to immune checkpoint inhibitor (ICI) therapy. To explore the potential therapeutic benefits of sulindac in immunotherapy, we treated two triple-negative breast cancer (TNBC) mouse models with low doses of sulindac, approximately equivalent to 1/8 of the FDA-approved human dose. Consistent with our previous findings, we found no significant differences in side effects among the treatment groups when assessing PGE2 levels in mouse blood samples. We also monitored the total body weight of the mice twice a week as a surrogate marker of toxicity. No significant body weight loss was observed following sulindac treatment. These results suggest that at a carefully controlled low dose, sulindac is safe and can enhance immune checkpoint inhibitor (ICI) therapy with minimal systemic toxicity.

Our conclusion is that sulindac effectively modulates TNBC’s response to anti-PD-L1 immunotherapy. Its mechanisms include downregulating PD-L1 by blocking Stat3 signaling and enhancing miR-570-3p expression, which potentially targets PD-L1. Given that patients with Triple-Negative Breast Cancer (TNBC) derive minimal benefit from current ICI regimens, our research offers unique insights into developing sulindac as a safe and effective component of a combined immunotherapy for treating TNBC.

## Acknowledgment

This work was supported in part by the National Institutes of Health (NIH)-National Cancer Institute (NCI) grants (R01CA271533, R01CA260698, and R01CA275089) to YX.

