## Supplemental figures for "Sulindac modulates the response of triple negative breast cancer to anti-PD-L1 immunotherapy"

**Supplementary Data:**

**Supplementary Figure 1:** The picture of 4T1 tumor

**Supplementary Figure 2:** Percentage of human CD45+ immune cells and human CD8+ T cells in the blood of PBMC humanized mice after PBMCs injection up to two weeks.

**Supplementary Figure 3:** The picture of 56S tumor.

**Supplementary Figure 4:** Average body weight of mice measured twice a week.

**Supplementary Figure 5:** Unmerged immunofluorescence images of CD8+ T cells in 4T1 tumor tissues. Green: CD8+ T cells; Blue: DAPI.

**Supplementary Figure 6:** Unmerged immunofluorescence images of CD8+ T cells in 56S tumor tissues. Green: CD8+ T cells; Blue: DAPI.

**Supplementary Figure 7:** Unmerged immunofluorescence images of CD8+ T cells in 56S tumor organoid. Green: CD8+ T cells; Blue: DAPI.

**Supplementary Figure 8:** Unmerged immunofluorescence images of CD8+ T cells in 56S tumor organoid mixed with activated human PBMCs and treated with vehicle control DMSO at 0.1%, 20 μM sulindac sulfide (SS), 200 nM anti-human PD-L1 antibody (Atezolizumab), and combination of 20 μM SS with 200 nM anti-human PD-L1 antibody (Atezolizumab) for 2 days. Green: CD8+ T cells; Blue: DAPI.

**Supplementary Figure 9:** Unmerged immunofluorescence images of PD-L1 and CD8+ T cells in 56S tumor tissues. Red: PD-L1; Green: CD8+ T cells; Blue: DAPI.

**Supplementary Figure 10:** Unmerged immunofluorescence images of PD-L1 and CD8+ T cells in 4T1 tumor tissues. Red: PD-L1; Green: CD8+ T cells; Blue: DAPI.

**Supplementary Figure 11:** Relative quantitation of exosomal PD-L1. Exomsome lysates from mouse plasma samples were incubated overnight with mouse PD-L1 antibody, CD63 antibody, and Calnexin antibody, respectively. CD63 and Calnexin are the positive and negative markers for identifying exosomes. 4T1 cell lysates were used as positive control for Calnexin antibody. Quantity One software (Bio-Rad) was used to calculate the intensity ratio between PD-L1 and CD63. The number of exosome particles were used to normalize the expression of mouse exosomal PD-L1.

**Supplementary Figure 12:** Relative quantitation of exosomal PD-L1. Exomsome lysates from mouse plasma samples were incubated overnight with human PD-L1 antibody, CD9 antibody, and Calnexin antibody, respectively. CD9 and Calnexin are the positive and negative markers for identifying exosomes. MDA-MB-231 cell lysates were used as positive control for Calnexin antibody. Quantity One software (Bio-Rad) was used to calculate the intensity ratio between PD-L1 and CD63. The number of exosome particles were used to normalize the expression of human exosomal PD-L1.

**Supplementary Figure 13:** (a-d) IL6 can induce the relative luciferase activity in 293Tn and MDA-MB-231 cells transfected with pGL3 luciferase reporter vector containing the sequence of wild type PD-L1 promoter. After transfection, cells were treated with IL6 for 4 h, and then the luciferase activity was measured using a dual luciferase reporter assay. RLU: relative luminescence units; t-test was used to determine statistical significance; **p<0.01; ***p<0.001

**Supplementary Figure 14:** (a-d) Stat3 pcDNA3 vector can induce the relative luciferase activity in 293Tn and MDA-MB-231cells transfected with pGL3 luciferase reporter vector containing the sequence of wild type PD-L1 promoter. The luciferase activity was measured using a dual luciferase reporter assay. RLU: relative luminescence units; t-test was used to determine statistical significance; ***p<0.001

**Supplementary Figure 15:** Unmerged immunofluorescence images of PD-L1 and p-Stat3 in E0771 cells. Red: p-Stat3; Green: PD-L1; Blue: DAPI.

**Supplementary Figure 16:** Unmerged immunofluorescence images of PD-L1 and p-Stat3 in 4T1 cells. Red: p-Stat3; Green: PD-L1; Blue: DAPI.

**Supplementary Figure 17:** Unmerged immunofluorescence images of PD-L1 and p-Stat3 in MDA-MB-231 cells. Red: p-Stat3; Green: PD-L1; Blue: DAPI.

**Supplementary Figure 18:** Unmerged immunofluorescence images of PD-L1 and p-Stat3 in HCC70 cells. Red: p-Stat3; Green: PD-L1; Blue: DAPI.

**Supplementary Figure 19:** Unmerged immunofluorescence images of PD-L1 and p-Stat3 in 4T1 tumor tissue. Red: p-Stat3; Green: PD-L1; Blue: DAPI.

**Supplementary Figure 20:** Unmerged immunofluorescence images of PD-L1 and p-Stat3 in 56S tumor tissue. Red: p-Stat3; Green: PD-L1; Blue: DAPI.

### Supplementary Figure 1

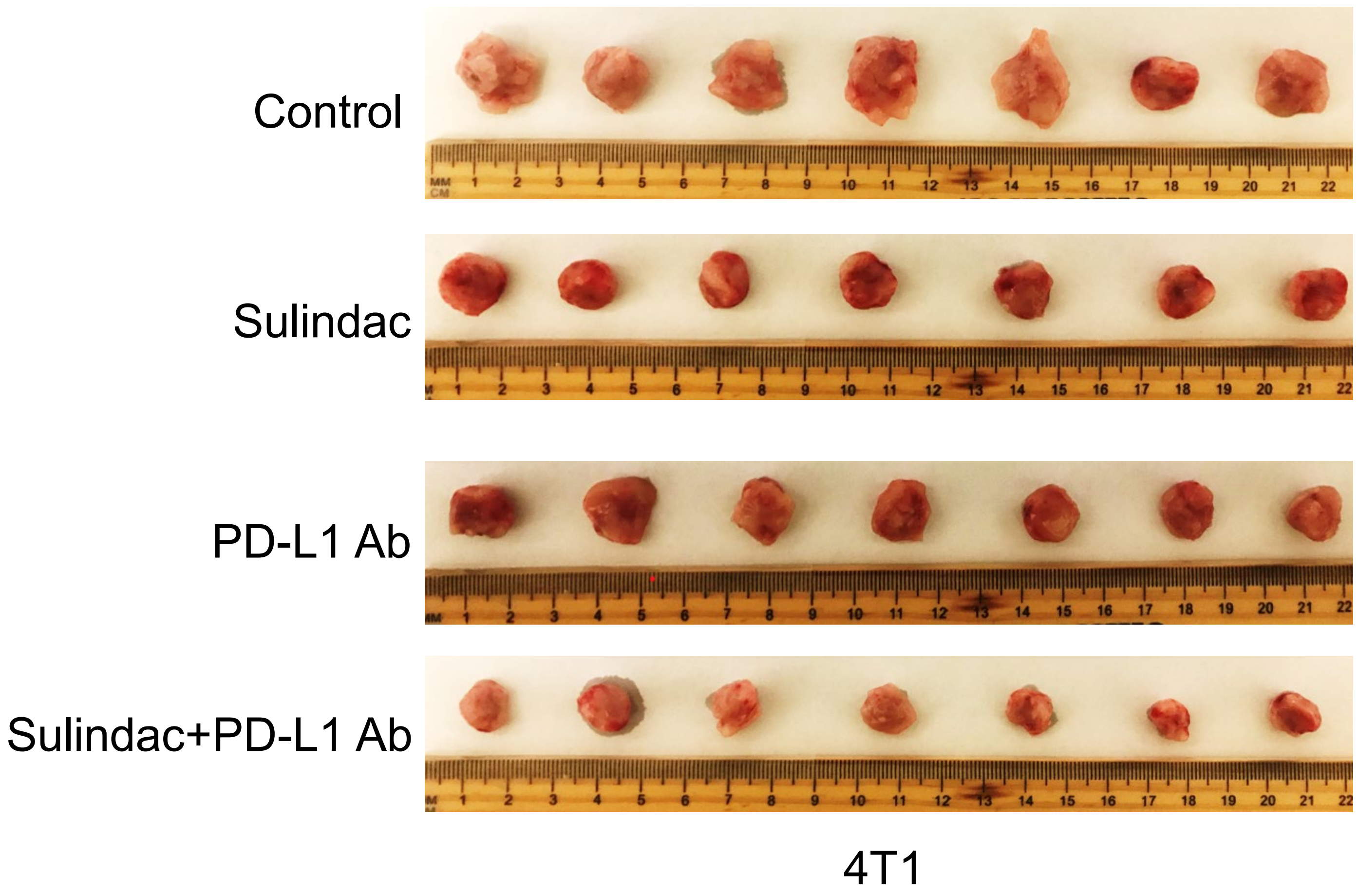

### Supplementary Figure 2

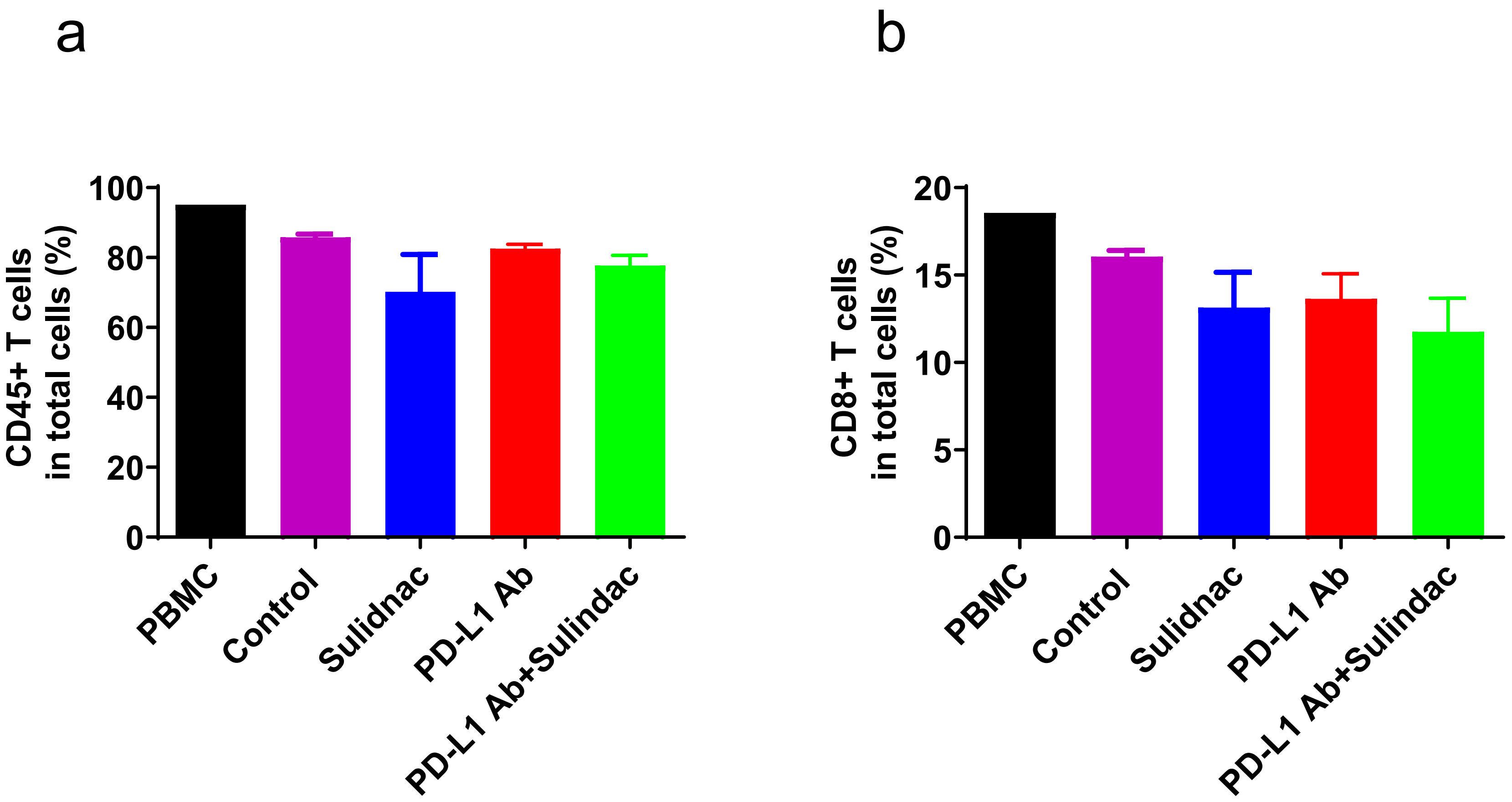

#### Supplementary Figure 3

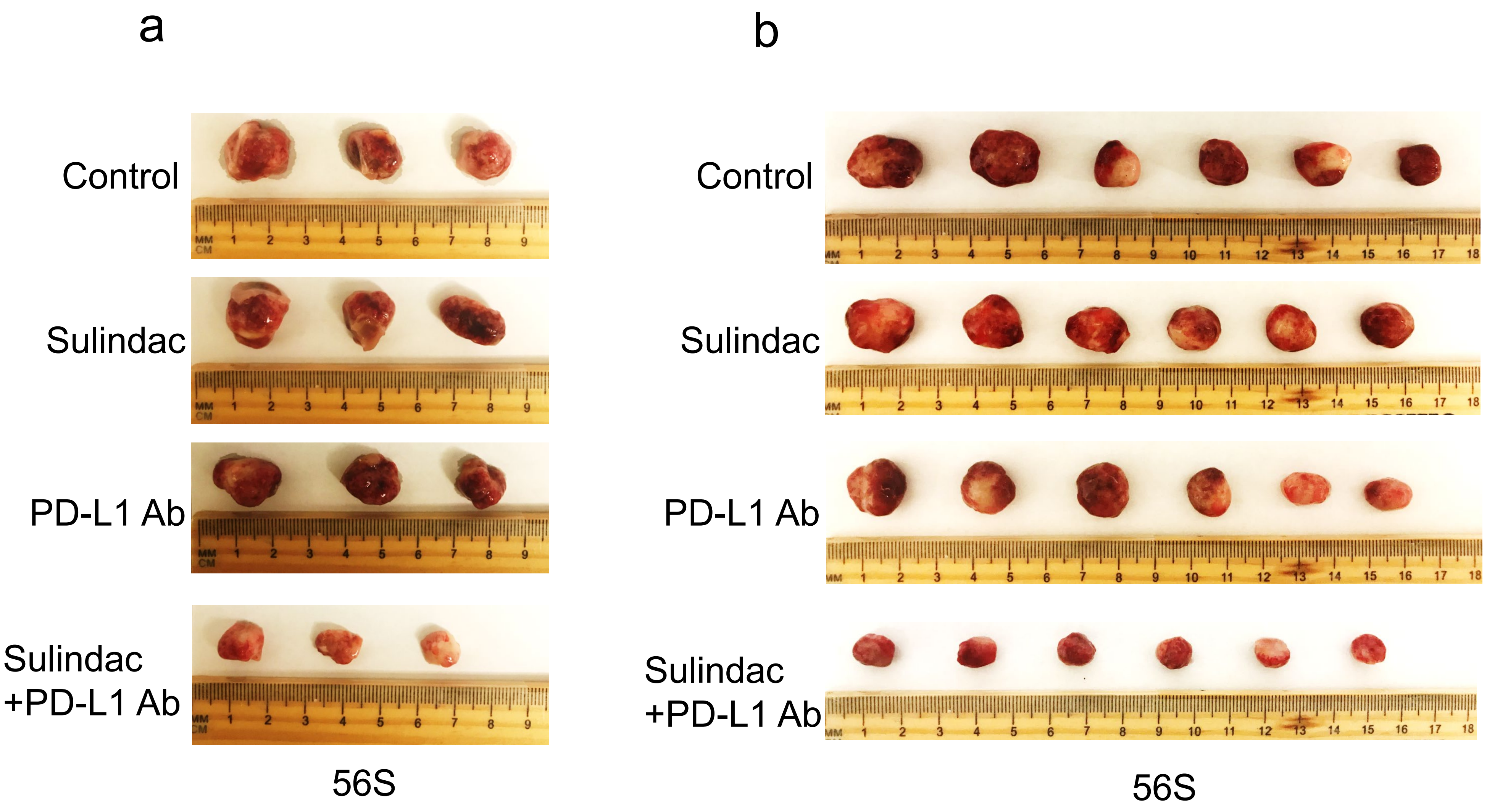

### Supplementary Figure 4

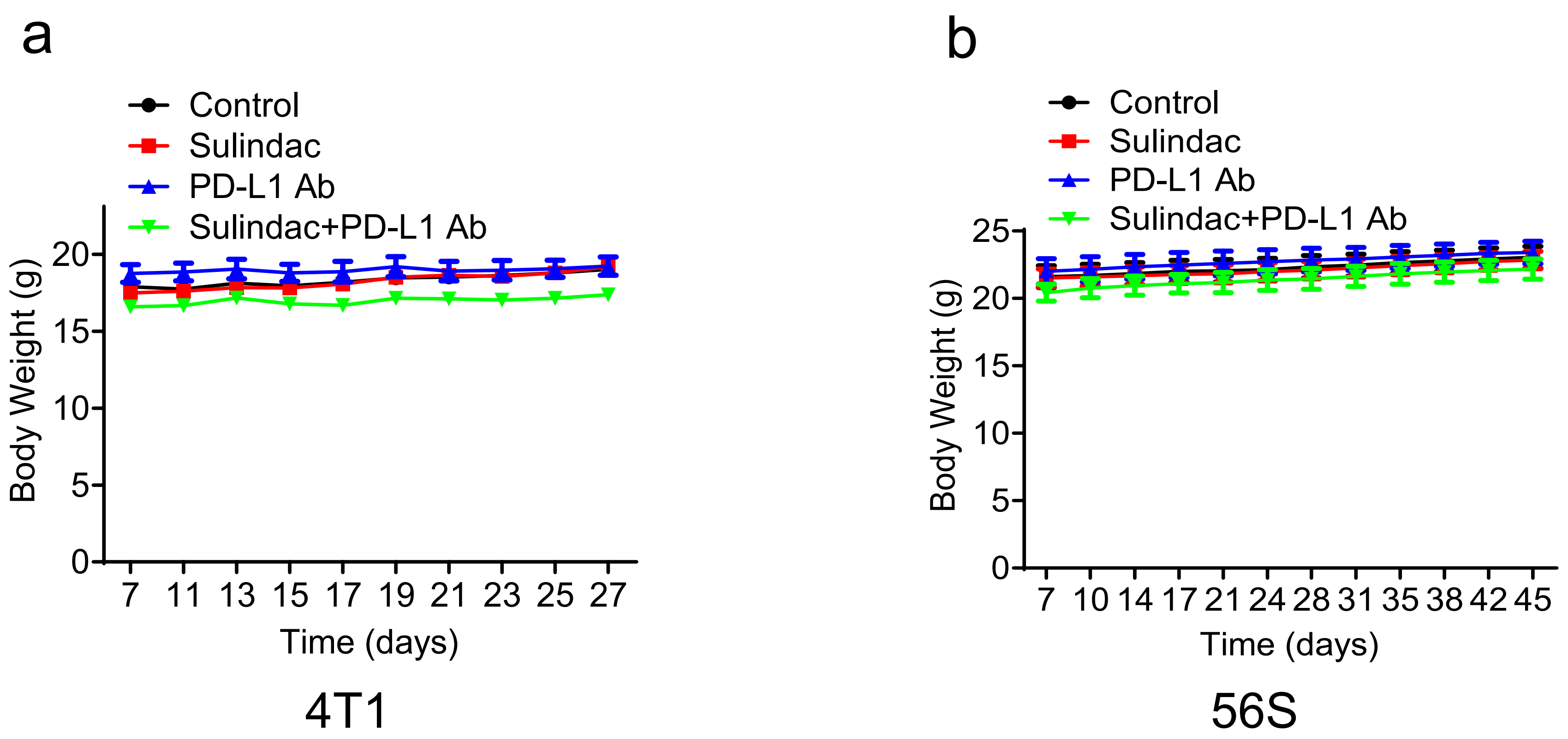

### Supplementary Figure 5

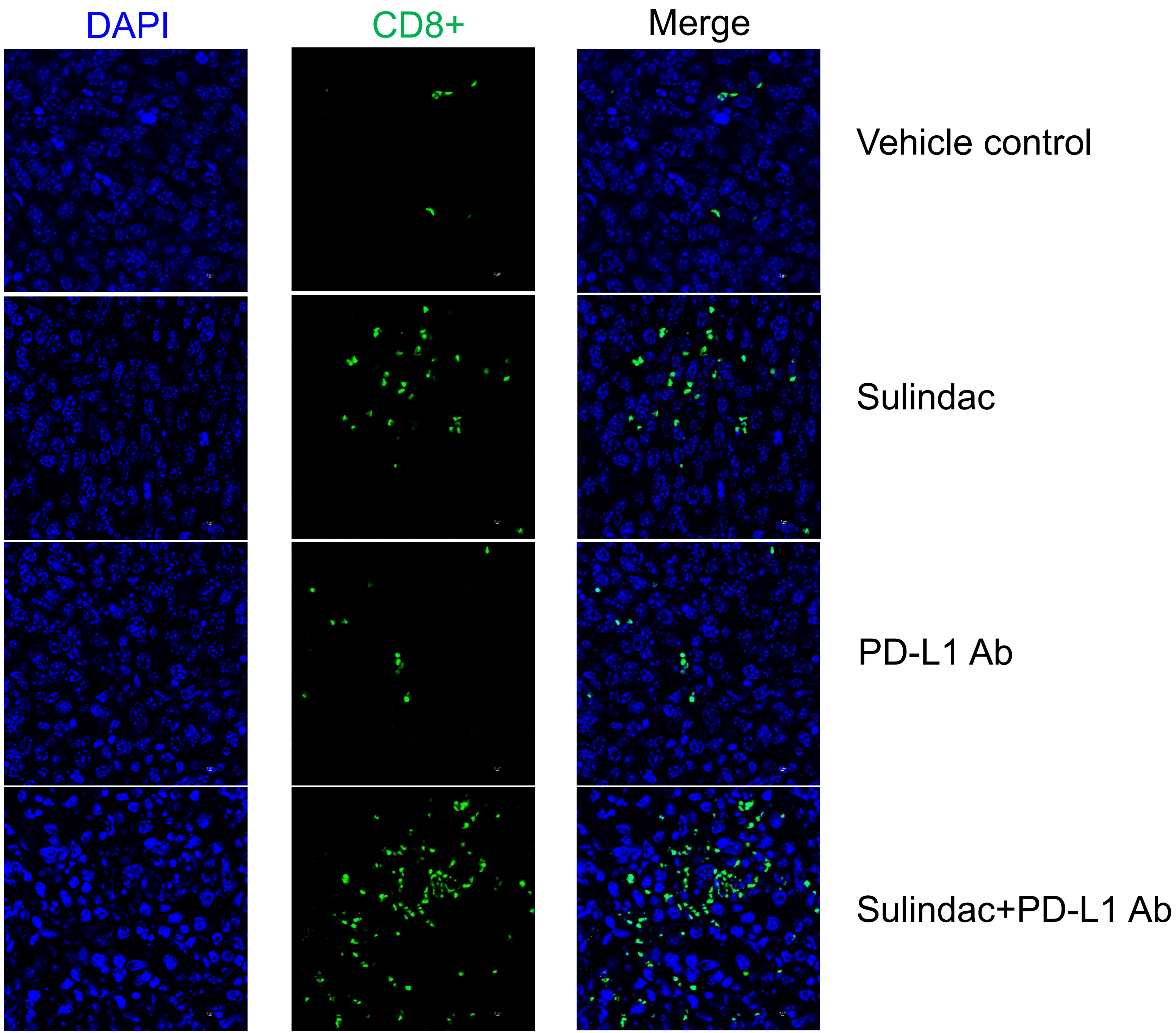


4T1 tumor tissue

### Supplementary Figure 6

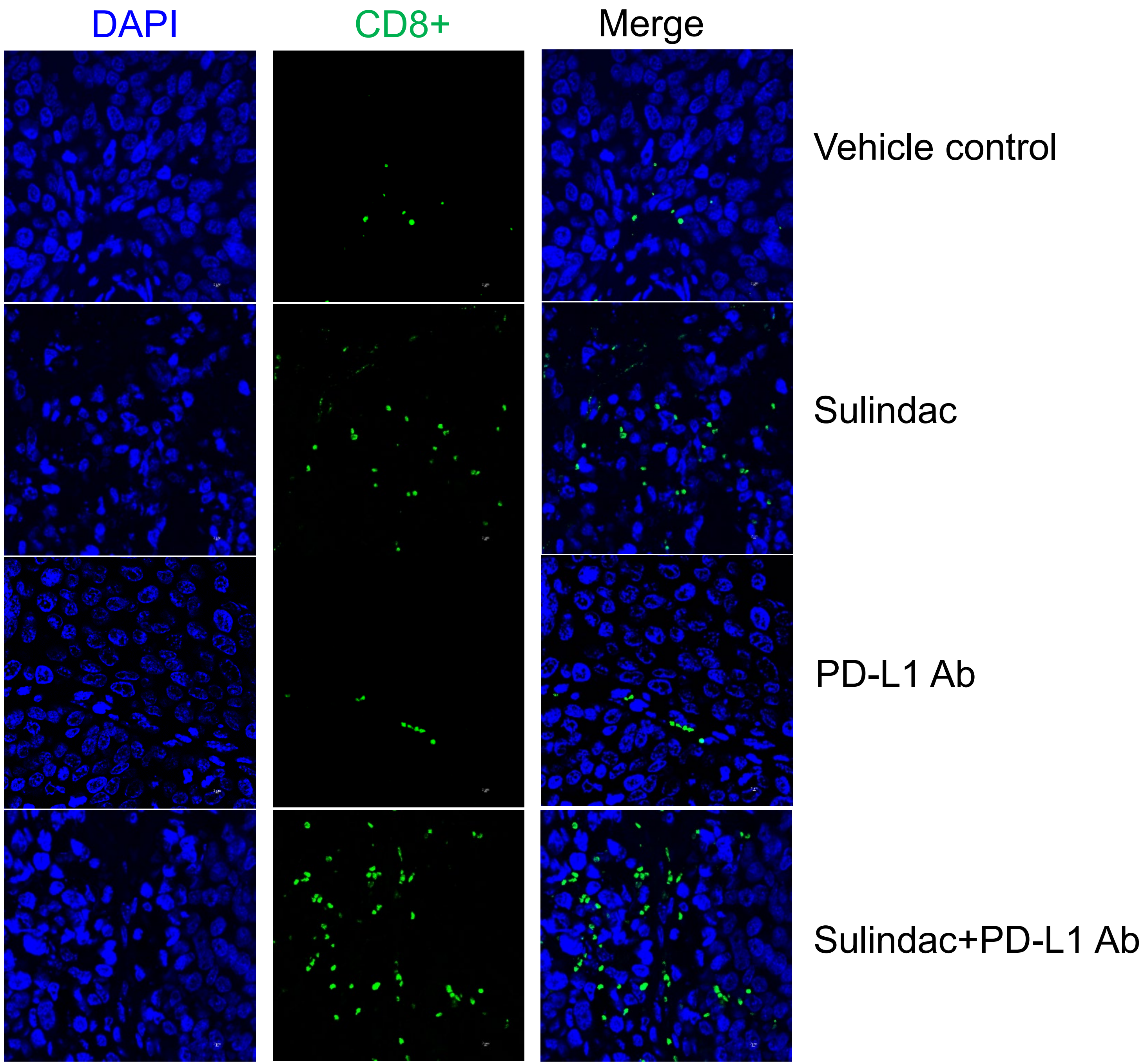


56S tumor tissue

### Supplementary Figure 7

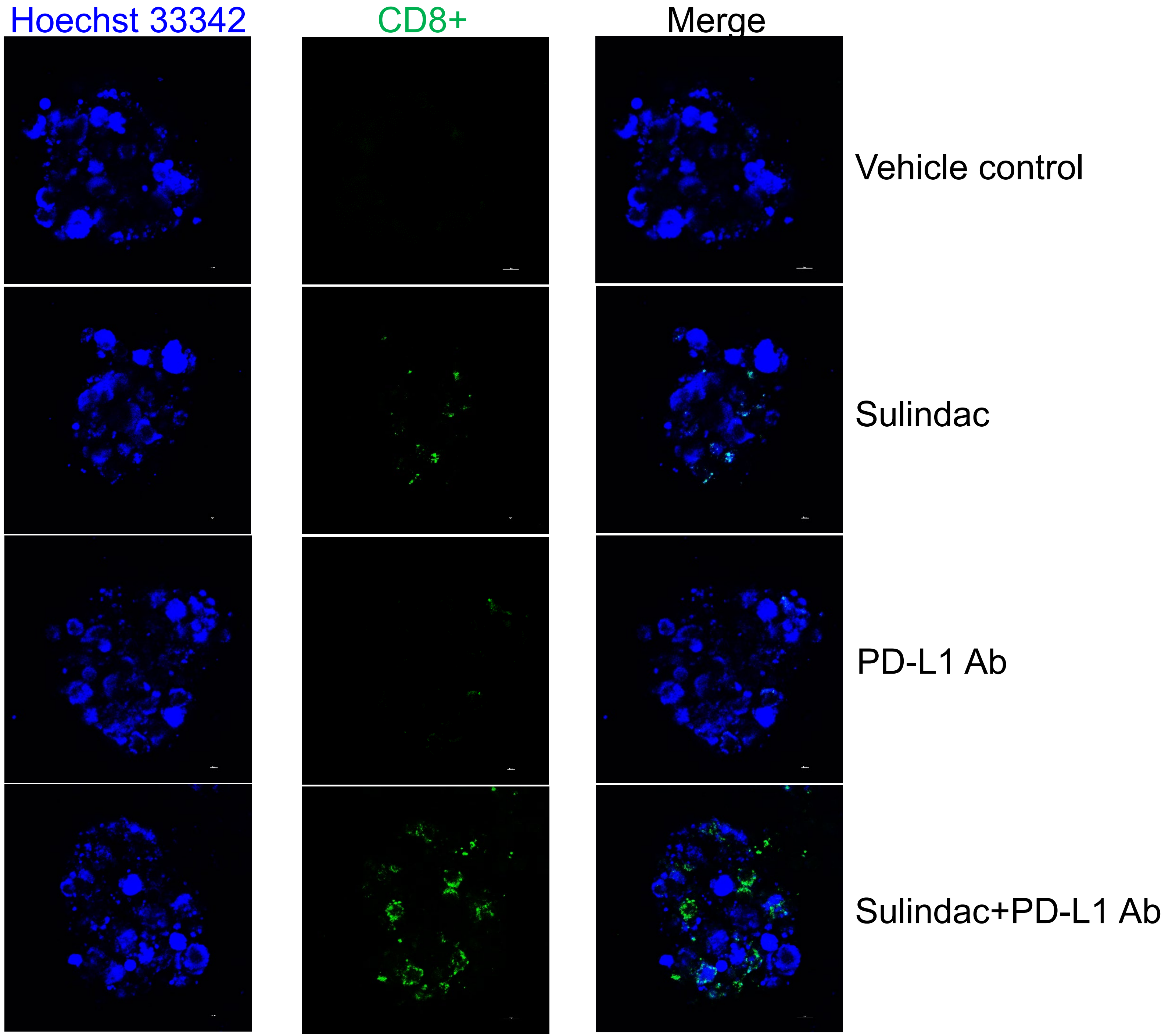


56S organoid

### Supplementary Figure 8

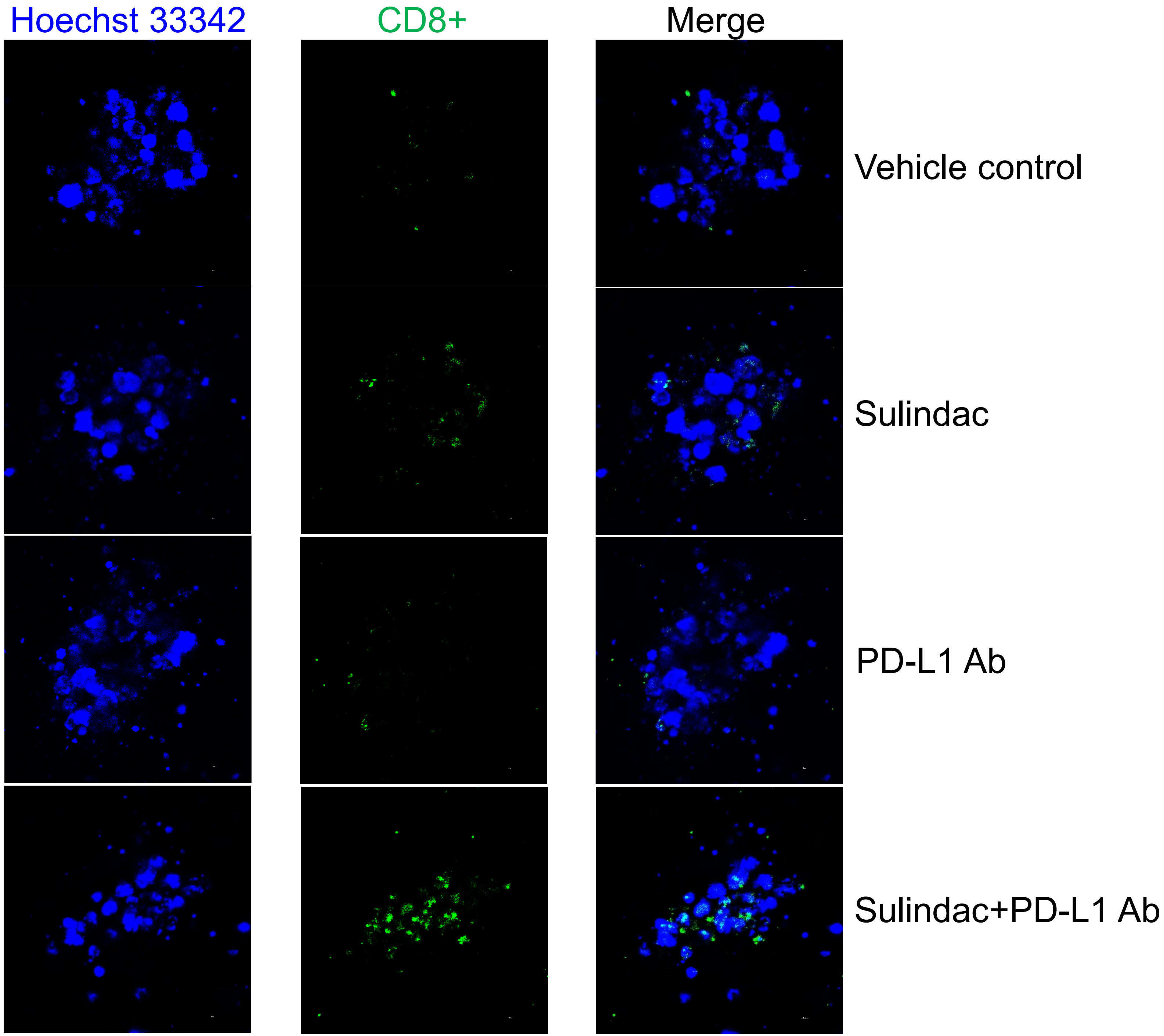


56S organoid

### Supplementary Figure 9

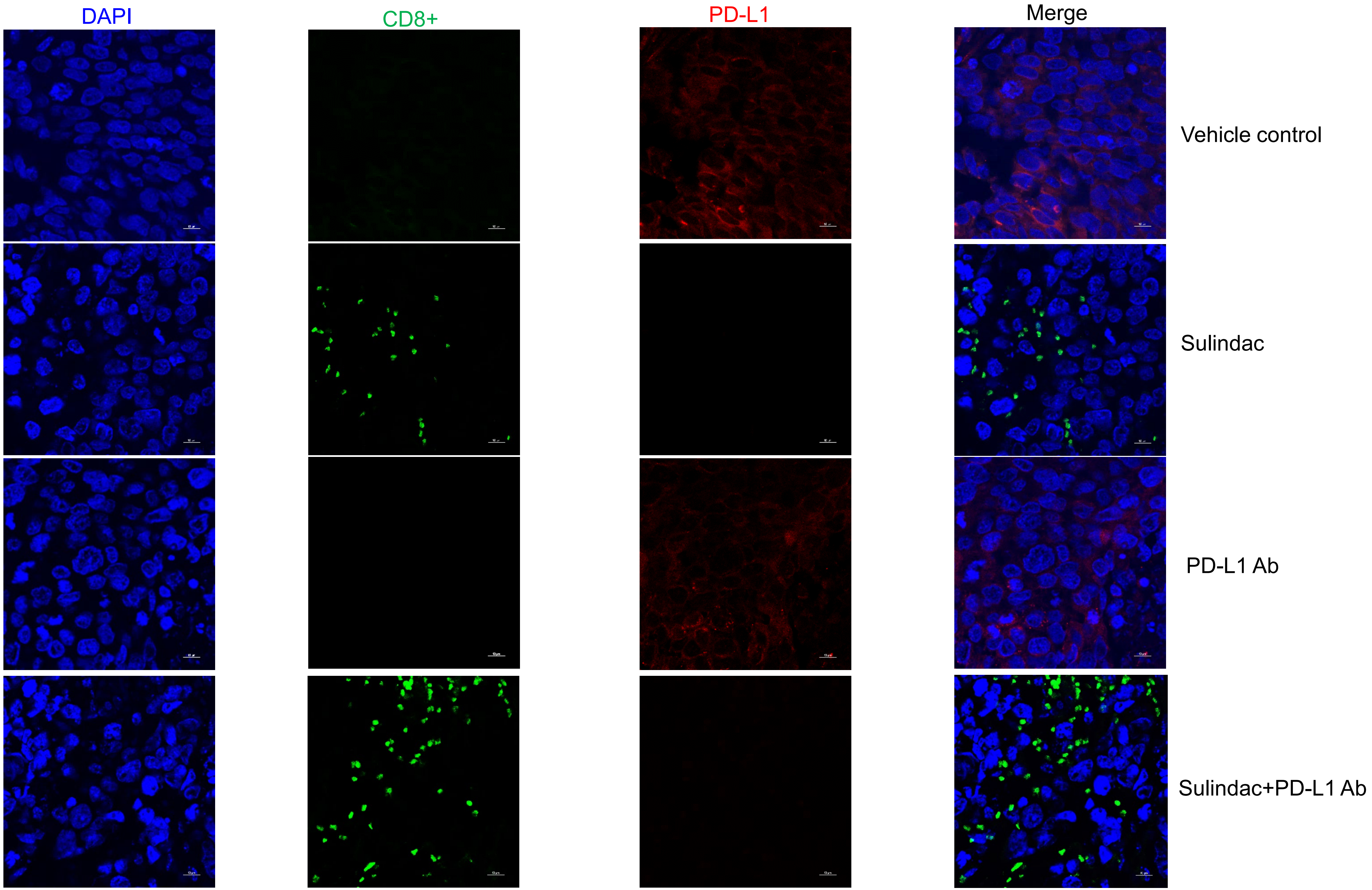


56S tumor tissue

### Supplementary Figure 10

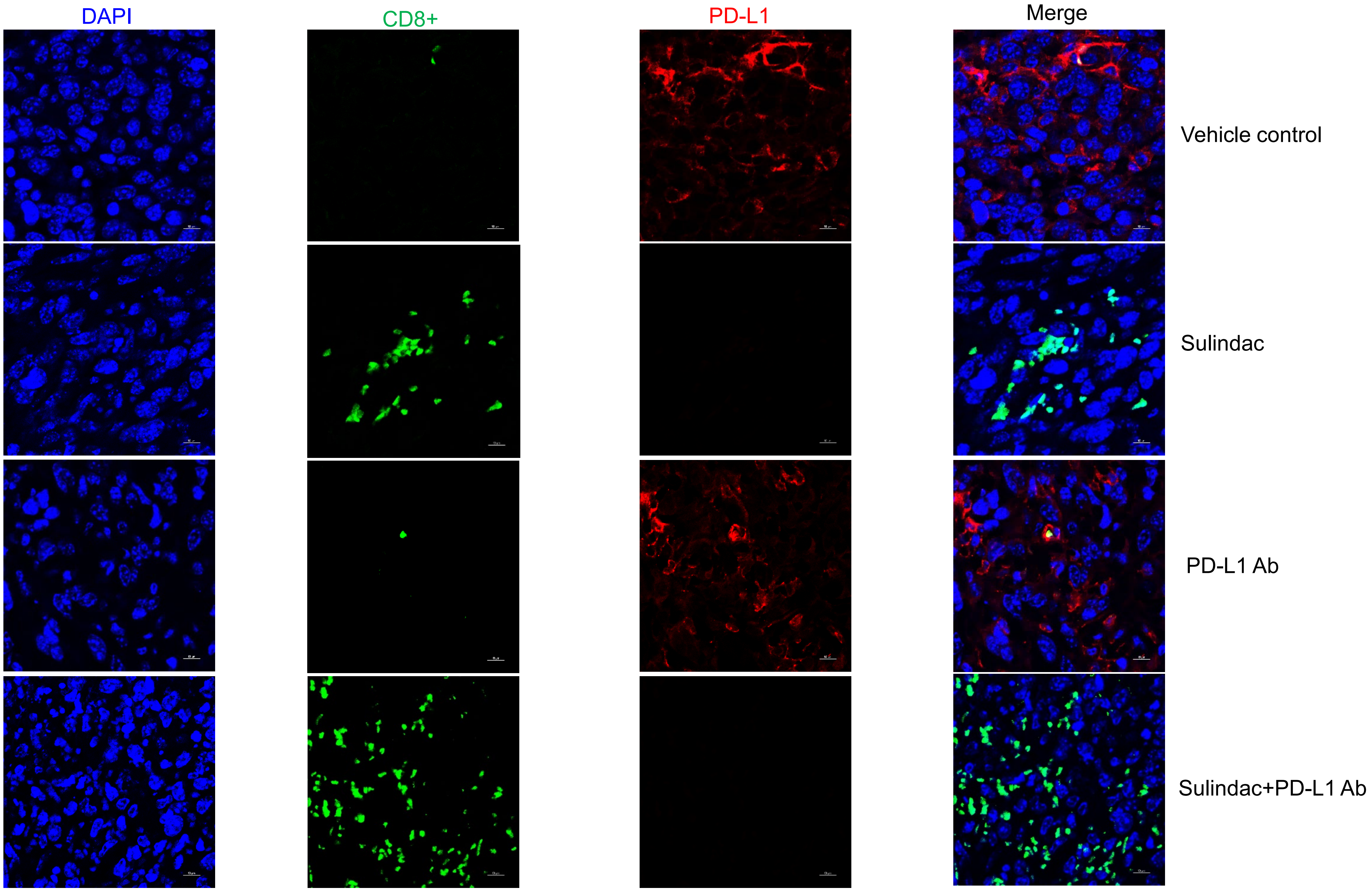


4T1 tumor tissue

### Supplementary Figure 11

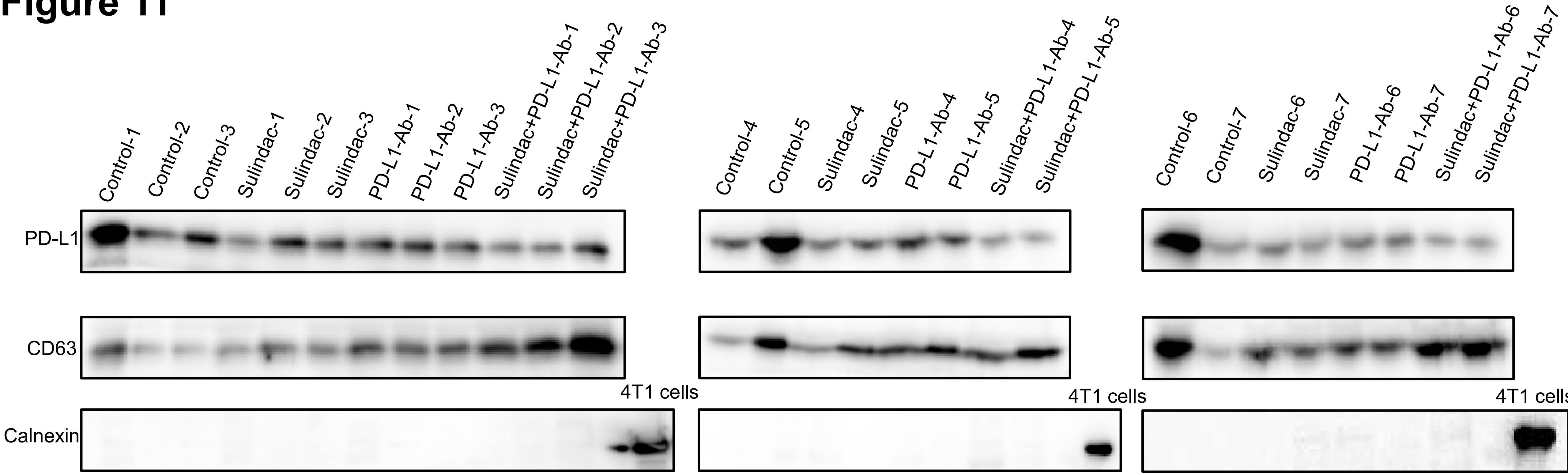

### Supplementary Figure 12

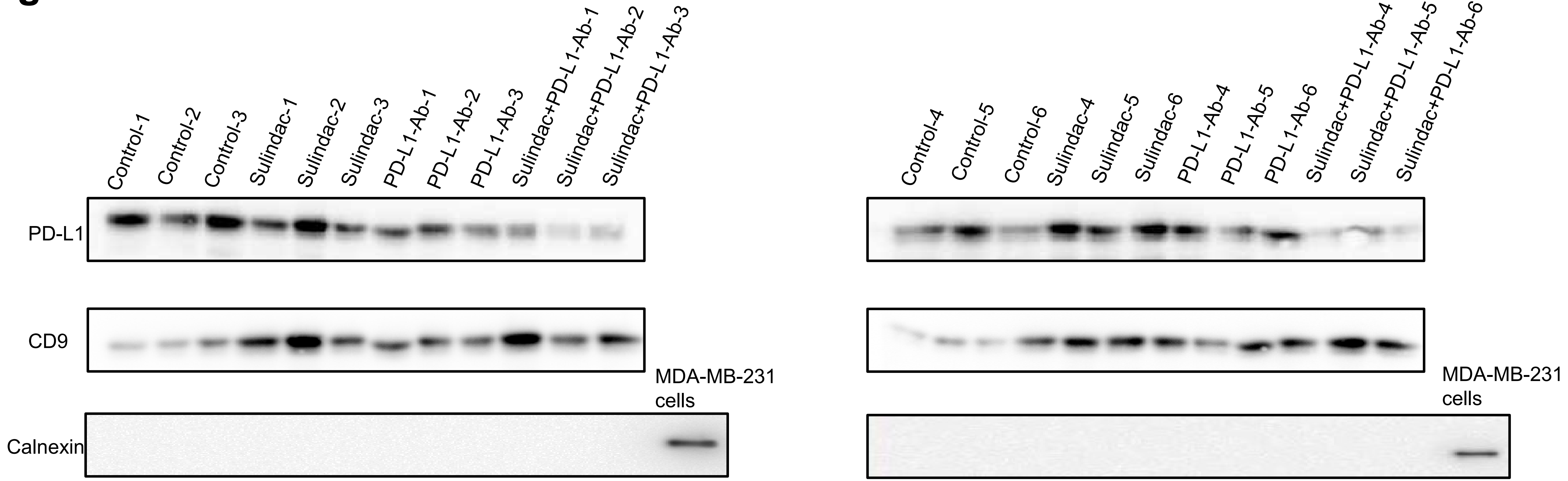

**Supplementary Figure 13**

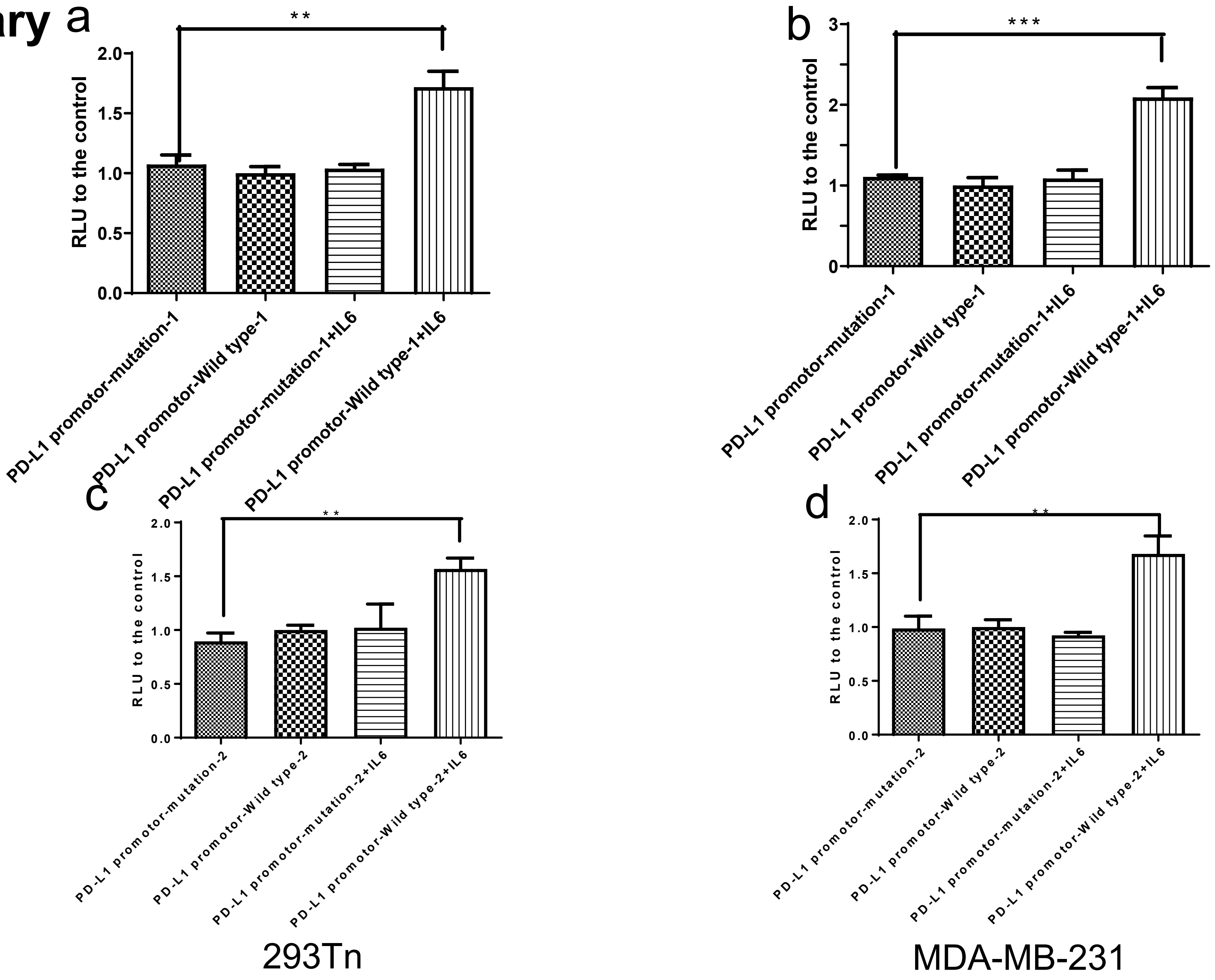

### Supplementary Figure 14

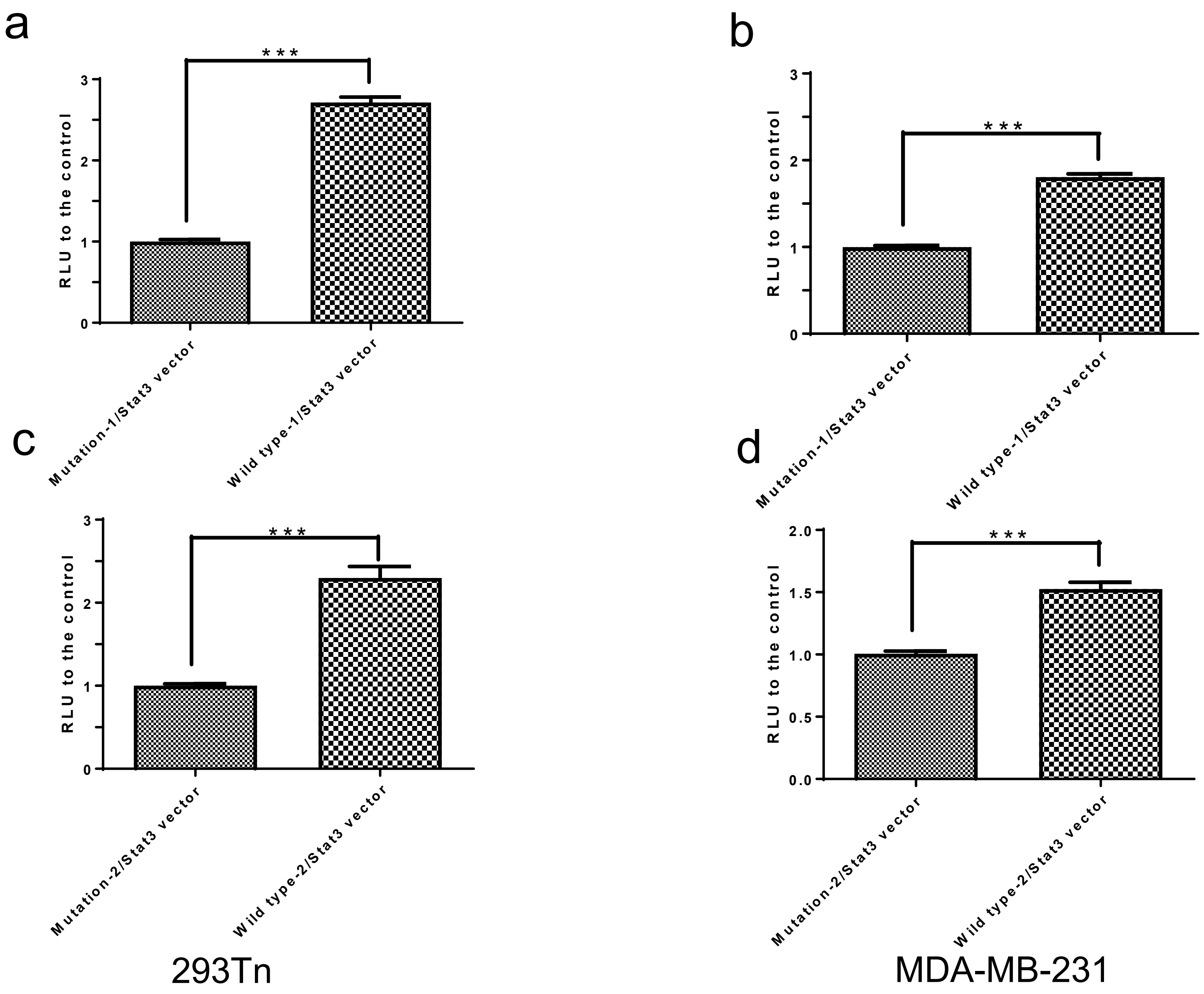

**Supplementary Figure 15**

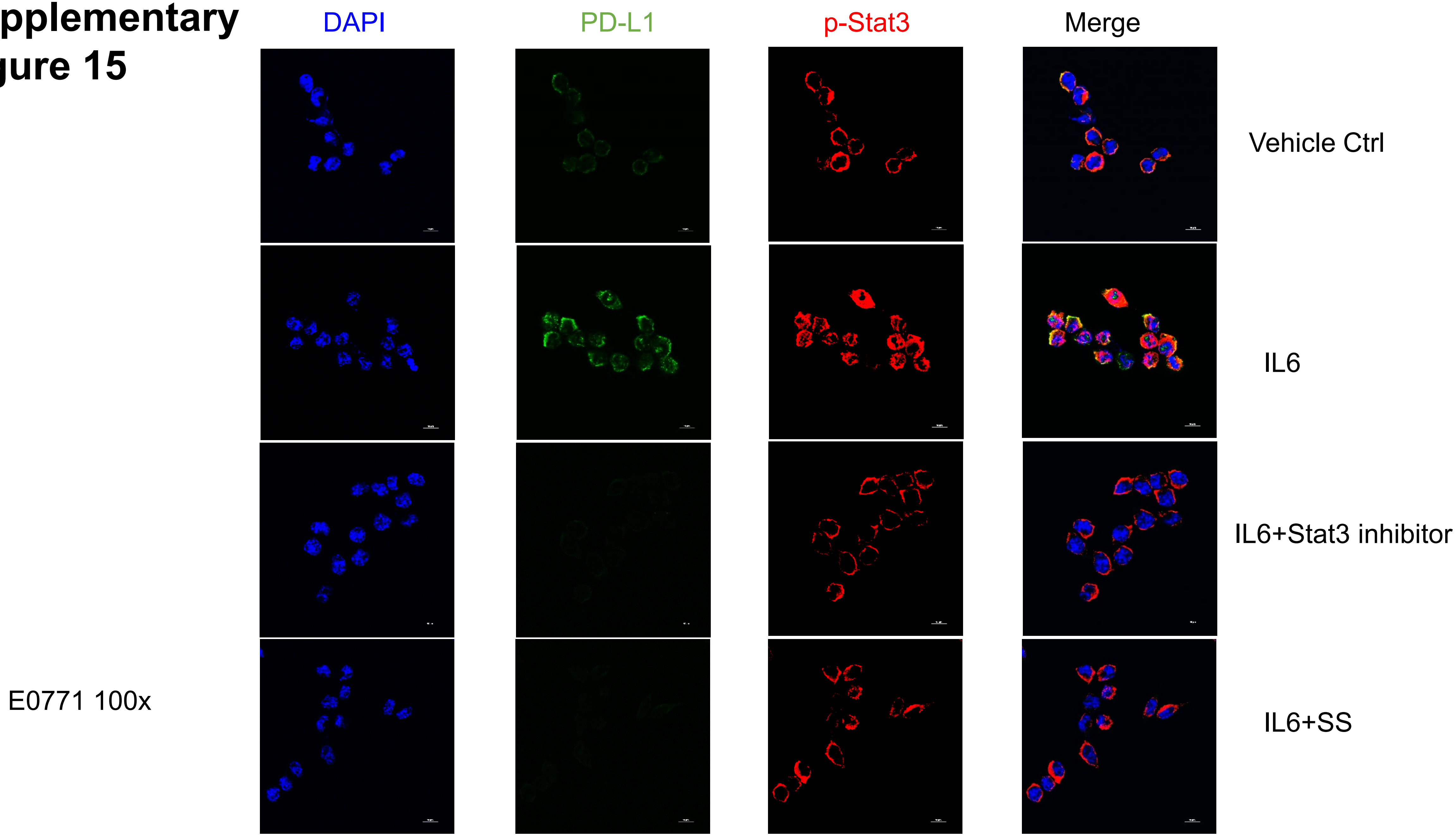

**Supplementary Figure 16**

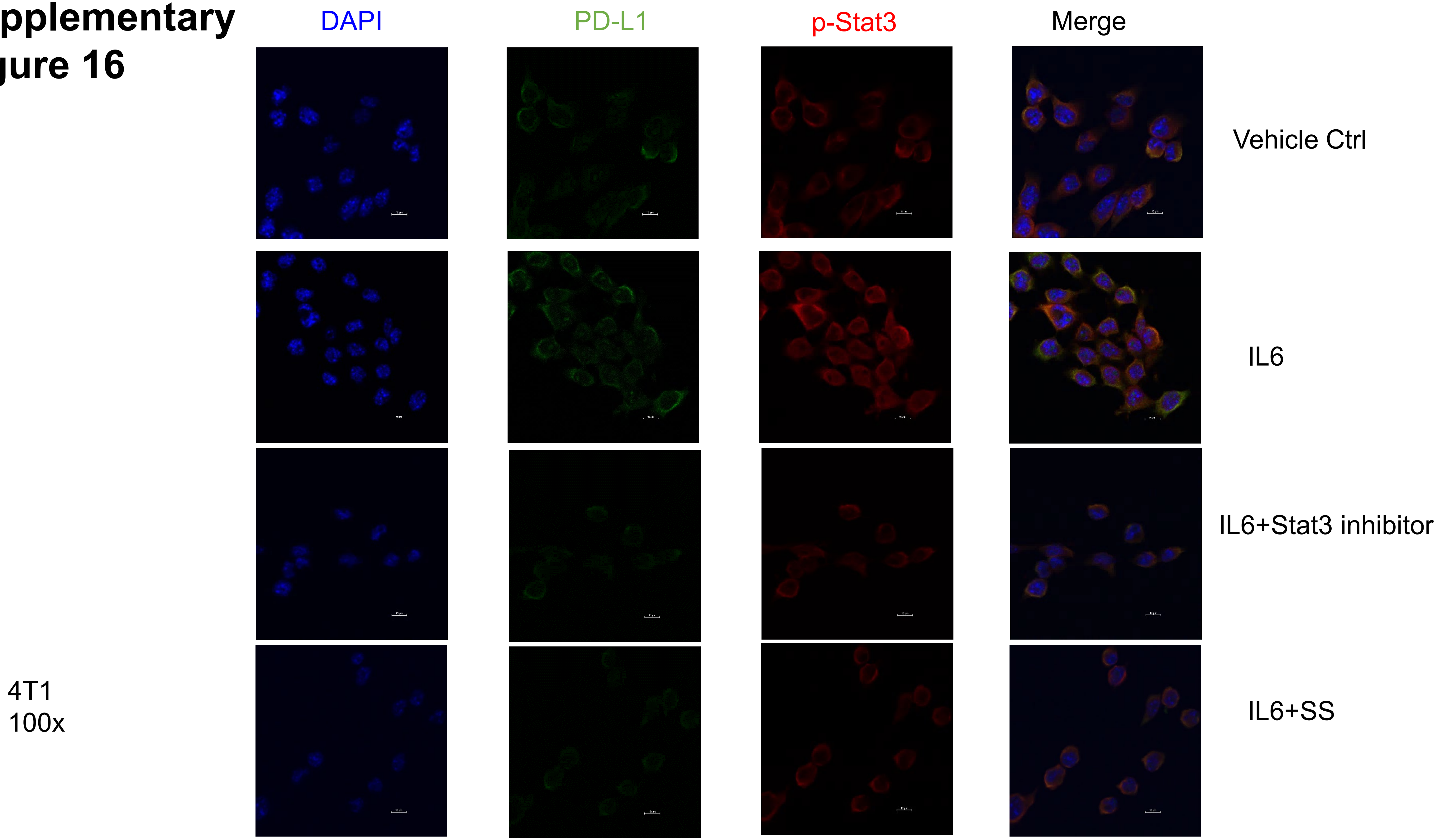

### Supplementary Figure 17

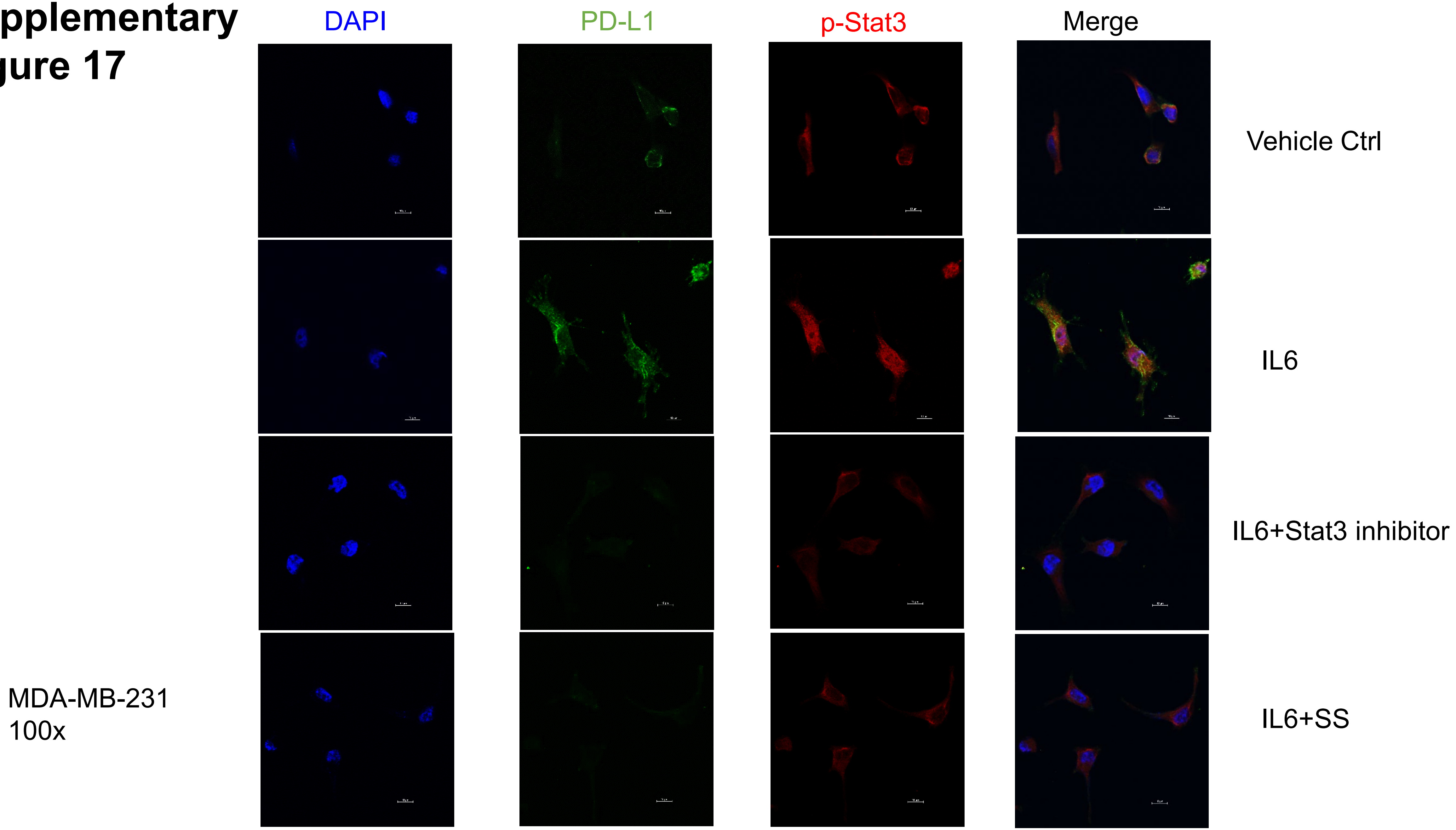

### Supplementary Figure 18

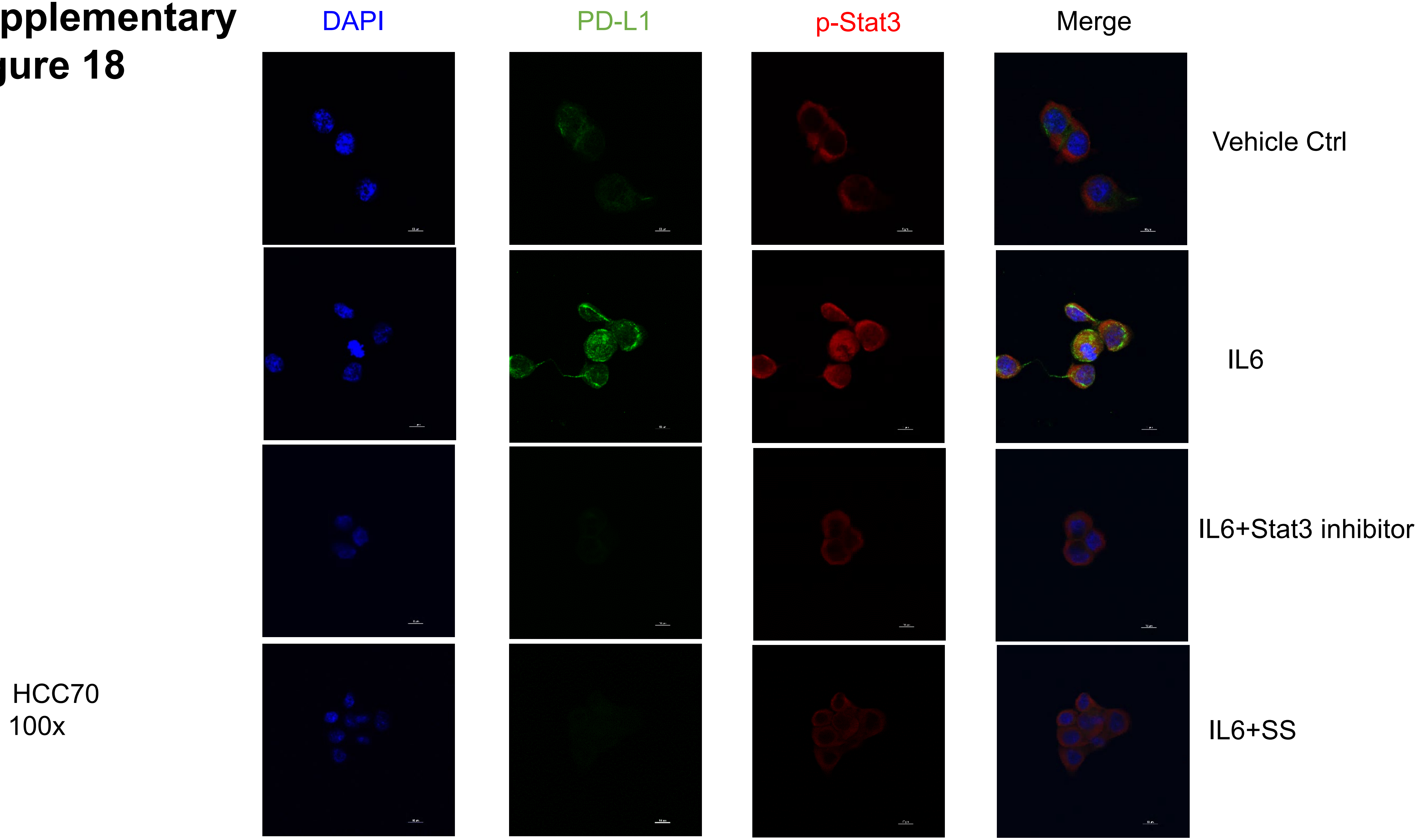

### Supplementary Figure 19

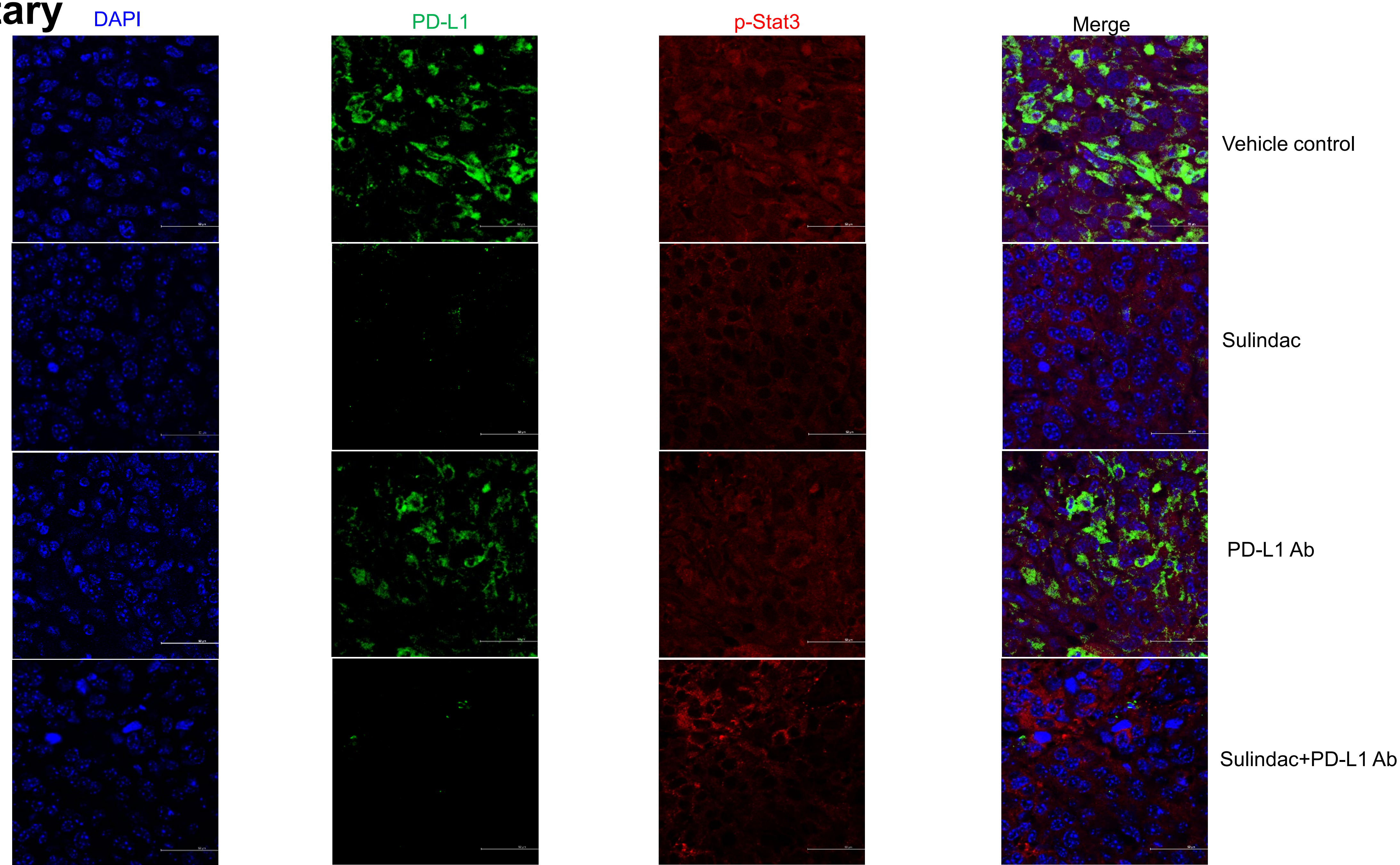


4T1 tumor tissue

### Supplementary Figure 20

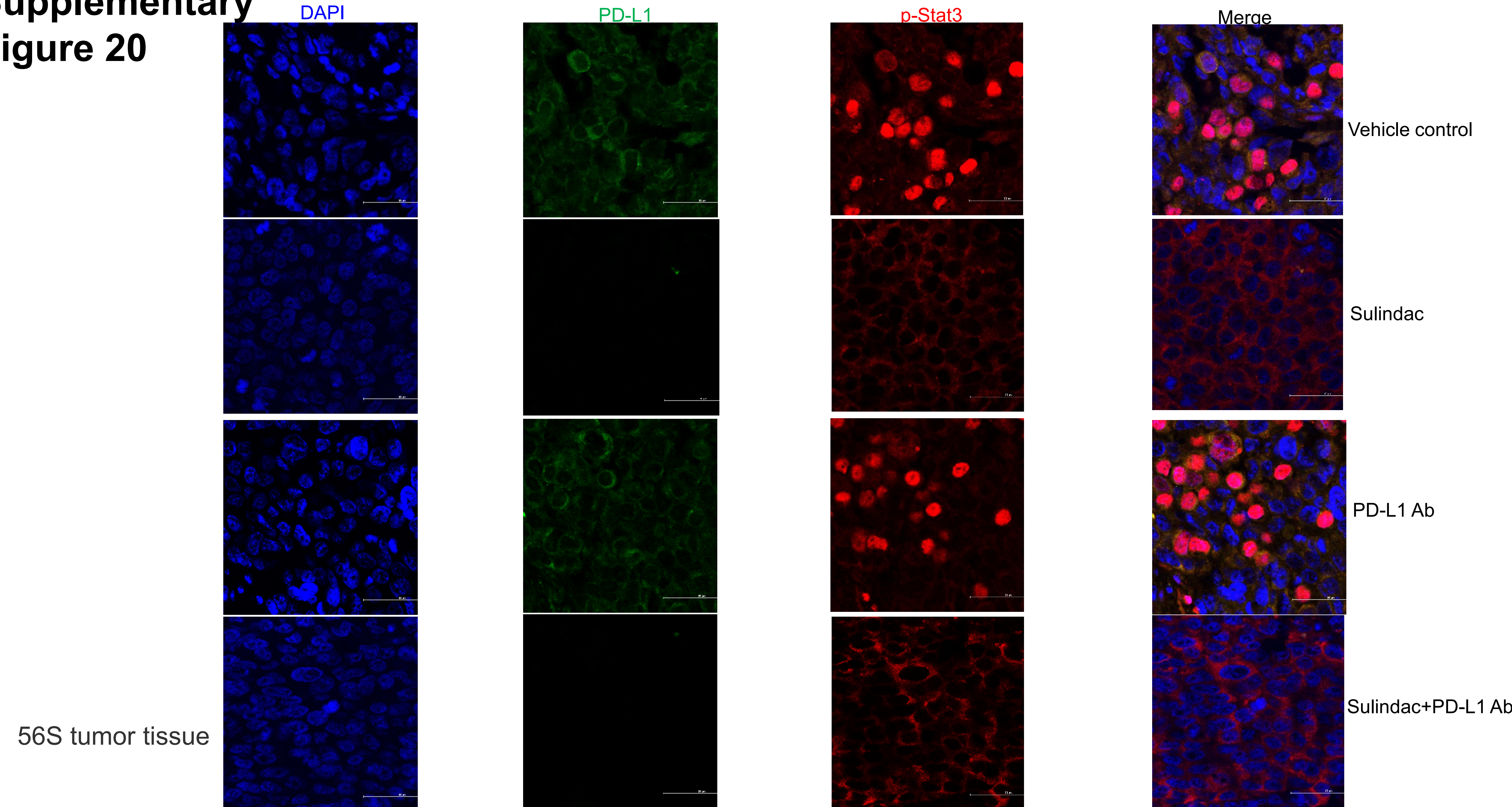
